# Ophiostomatoid fungi associated with bark beetles infesting *Pinus armandii*, with characterizations of seven new species from China

**DOI:** 10.1101/2024.11.06.622300

**Authors:** Minjie Chen, Yutong Ran, Congwang Liu, Kun Liu, Tong Lin, Mingliang Yin

## Abstract

Ophiostomatoid fungi mainly refer to the order Ophiostomatales (Ascomycota), which are known to be symbiotic with bark beetles infesting pine trees. These fungi are primarily associated with *Dendroctonus armandi* and other bark beetles, significantly impacting the local pines *Pinus armandii* in western China. However, this fungal diversity must be further explored. This study aims to identify Ophiostomatoid fungi associated with *P. armandii* in Ningshaan, Shaanxi Province. A total of 225 Ophiostomatalean isolates were obtained from 46 gallery samples, accounting for 77.6 % of the total number of isolated strains. Twenty species were identified based on the phylogenetic analyses of ITS, LSU, β-tubulin, and EF-1α, as well as morphological characteristics. These include seven new taxa, *Ceratocystiopsis rugosa*, *Graphilbum prolificum*, *G. ramosus*, *Masuyamyces huoditangensis*, *Ophiostoma flavescens*, *O. lenta*, *O. mediotumida*, and thirteen known species. This study laid a theoretical foundation for understanding the relationship between the beetles and the ophiostomatalean fungi in the Ningshan region, China.

## INTRODUCTION

Bark beetles represent a remarkably diverse group of insects characterized by their phloem-feeding behavior and complex evolutionary relationships with various symbiotic organisms. Among these symbionts, a particularly significant group is the ophiostomatoid fungi, which are named for their distinctive morphological characteristics (De Beer et al., 2013). This taxonomic diversity reflects the complex evolutionary history of these fungal symbionts and their adaptation to various ecological niches associated with bark beetles. It not only provides the necessary nutrients for the life cycle of beetles but also improves the survival rate (Ayres et al. 2000; Lee et al. 2005; Kirisits et al. 2007; Wadke et al. 2016; Kandasamy et al. 2019). It is the key to successfully colonizing bark beetles on host trees (Wingfield et al. 1993; Kirisits et al. 2007; Diguistini et al. 2011; Zaman et al. 2023). In the symbiotic relationship between ophiostomatoid fungi and bark beetles carry spores through exoskeletons or actively transport and cultivate spores to inoculate their fungal partners (Raffa et al. 2015). There are also some special structural traps or depressions on the surface of the small brood beetles, which can help the fungus carry spores for transmission (Batra, 1963; Hofstetter et al., 2015; Kostovcik et al., 2015).

Ophiostomatoid fungi are a complex group of wood-infected fungi. Most species can grow and reproduce rapidly under suitable conditions. A large number of mycelia gather together to destroy their dredging system so that host plants cannot absorb soil moisture and cannot diffuse their nutrients, resulting in final death. Some pathogens even directly produce many toxic substances, which directly make the host trees lose their ability to live (Huang et al. 2015). Well-known examples include *Ophiostoma ulmi* and *O. novo-ulmi,* causing the Dutch elm wilt disease (De Hoog, 1974; Brasier, 1991), black root disease caused by *Leptographium wageneri* (Harrington et al. 1988), and laurel wilt disease caused by *Raffaelea lauricola* (Harrington et al. 2008). With the impact of factors such as global trade and climate change, the synergy of microbial pathogens and vectors has caused forest decline worldwide and posed a considerable threat to global biosafety. Therefore, it is urgent to explore the relationship between vector-fungus-host (Kenneth et al. 2008; Breda et al. 2014; Wingfield et al. 2015).

Zipfel et al. (2006) reconsidered the phylogenetic relationship of 50 species of *Ophiostoma* based on the DNA sequence numbers of LSU and β-tubulin genes, proved that *Ophiostoma* and *Grosmannia* were two independent genera, and redefined *Ceratocystiopsis*. De Beer et al. (2013) combined and reanalyzed the DNA sequences of the LSU and SSU of several research data and redefined the Ophiostomatales and its only family by constructing a phylogenetic tree. Based on phylogenetic analyses, Nel et al. (2021) introduced two new genera, *Intubia* and *Chrysosphaeria*. De Beer et al. (2022) recently conducted a phylogenetic analysis based on different DNA gene regions. The results confirm the division of nine genera among the currently recognized 14 genera, redefining *Grosmannia* and *Dryadomyces* (De Beer et al., 2022). With the deepening of research, more and more countries have extensively researched the diversity, pathogenicity, and interrelationships of Ophiostomatales, especially in North America, Europe, Japan, and China (Hulcr et al. 2020).

Pest damage in the northeast, southwest, northwest, and other regions severely endangers pine forest development in China. In the past ten years, there have been more and more reports on the diversity of associated ophiostomatoid fungi in the hosts infected by *E. vulgaris* in China. Still, the primary host plants are *Pinus koraiensis*, *Larix gmelinii*, *Pinus yunnanensis*, *Pinus cassia* and *Picea crassifolia*, etc., and there are few studies on the diversity of associated ophiostomatoid fungi in *P. armandii* (Yin et al. 2016; Chang et al. 2017; Chang et al. 2019; Chang et al. 2020; Pan et al. 2020; Wang et al. 2020; Wang et al. 2022). *Pinus armandii* is a native coniferous species and pioneer tree species in China. Since 1954, it has been continuously invaded by various bark beetles dominated by *Dendroctonus armandii*, resulting in many 20-30-year-old *Pinus armandii* deaths (Tang et al. 1999). Tang et al. (1999) found two unnamed fungi (*Ophiostoma* and *Leptographium* spp. ) associated with *Dendroctonus armandii.* They proved they were the triggers to overcome the host tree resistance system and the leading cause of its death. Tang et al. (2004) discovered a new companion fungus, *Leptographium qinlingensis*. Xie et al. (2008) found *Ceratocystis polonica*, a blue-stained fungus associated with *Dendroctonus armandii* of *Pinus armandii*, and inoculated it on healthy *Pinus armandii*. It was found to be an important pathogen that causes the host’s resin metabolism and water metabolism. Two ophiostomatoid fungi, *Ophiostoma brevicolle* and *Leptographium* sp. (Hu et al. 2015), were found in the study of the community structure of intestinal fungi at different developmental stages of *D. armandi*. In 2018, when studying the fungal symbionts of *E. minor* in China and Vietnam on Asian pine species, three associated ophiostomatoid fungi, *Leptographium* sp., *Ophiostoma quercus,* and *Ophiostoma floccosum*, were found on the body surface of *D. armandi* (Skelton et al., 2018). Wang et al. (2022) identified three known species, *Esteyea vermicola*, *G. pseudormiticum,* and *L. wushanense*, two new taxa, *G. parakesiyea* and *O. shennongense*, an undetermined *Ophiostoma* sp. and a new *L. qinlingense*.

This study conducted field sampling, laboratory isolation, and culture analysis on the fungal community associated with bark beetles in the *Pinus armandii* ecosystem in Shaanxi Province, China. The species were identified by morphological observation and multi-site DNA sequence phylogenetic analysis. The results of this study provide a new understanding of the diversity of ophiostomatoid fungi associated with bark beetles in Shaanxi Province and offer a scientific theoretical basis for the occurrence and management of bark beetles.

## MATERIALS AND METHODS

### Sample Collection and Fungi Isolation

In August 2022, samples, including beetle adults and their breeding galleries, were collected from *P. armandii* infected with beetles at Huoditang Forest Farm (33 ° 25 ′ N, 108 ° 25 ′ E). Chengguan Town (32 ° 32 ′ N, 107 ° 53 ′ E) in Ankang City, Shaanxi Province, western China (Fig.1). The collected beetle adults and their galleries were placed in sterile centrifuge tubes and sterile sampling bags, respectively. After labeling, they were transported back to the laboratory and stored at four °C until the fungal separation was completed. The fungi observed in the galleries and pupal chamber were inoculated into 2 % malt extract agar with a diameter of 90 mm (MEA: 20 g BioLab malt extract powder, 20 g BioLab agar, and 1000 ml deionized water) by stereomicroscope. After injection, they were cultured in a dark environment at 25 °C for one to two weeks, and then the hyphae at the tip of the colony were cut and inoculated onto MEA plates to obtain pure strains. After the preliminary analysis of the macroscopic and microscopic characteristics of the pure strains, the representative strains of each morphology were selected for in-depth morphological, physiological, and molecular studies. All fungal isolates were preserved in the South China Agricultural University, Guangzhou, Guangdong Province, China (SCAU; for accession numbers, see Table 1). Ex-type cultures of ophiostomatoid fungi described in this study are deposited in the China General Microbiological Culture Collection Center (CGMCC), and the Holotype specimens (dry cultures) were deposited in the Herbarium Mycologicum, Academiae Sinicae (HMAS), Beijing, China.

**Fig. 1.**
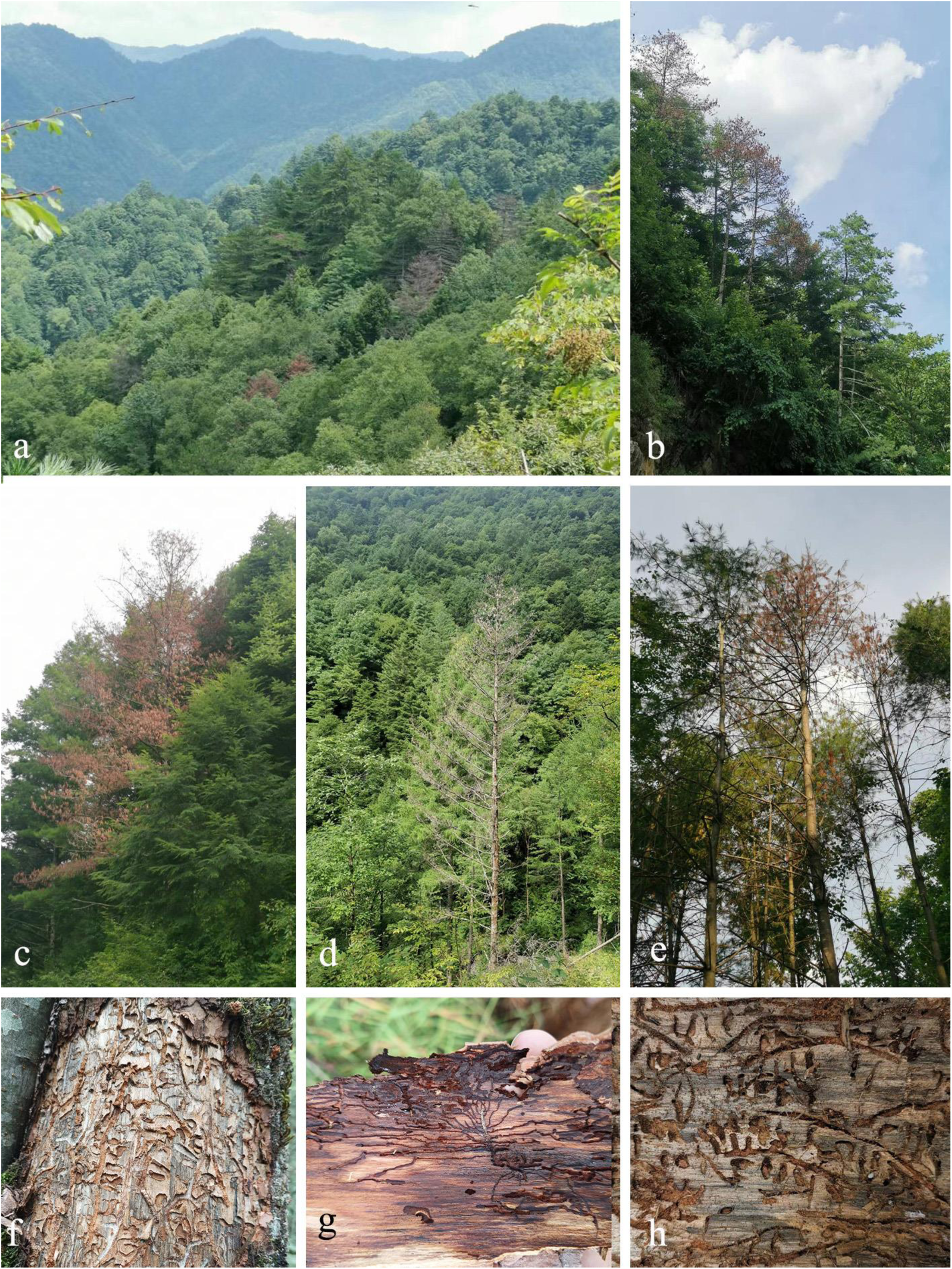
(a–e) Disease symptoms on *Pinus armandii* infested by bark beetles and ophiostomatoid fungi in western China; (f) Xylem after being eaten by bark beetles. (g-h) Galleries from *P. armandii* infested with bark beetles .

**Table 1.**
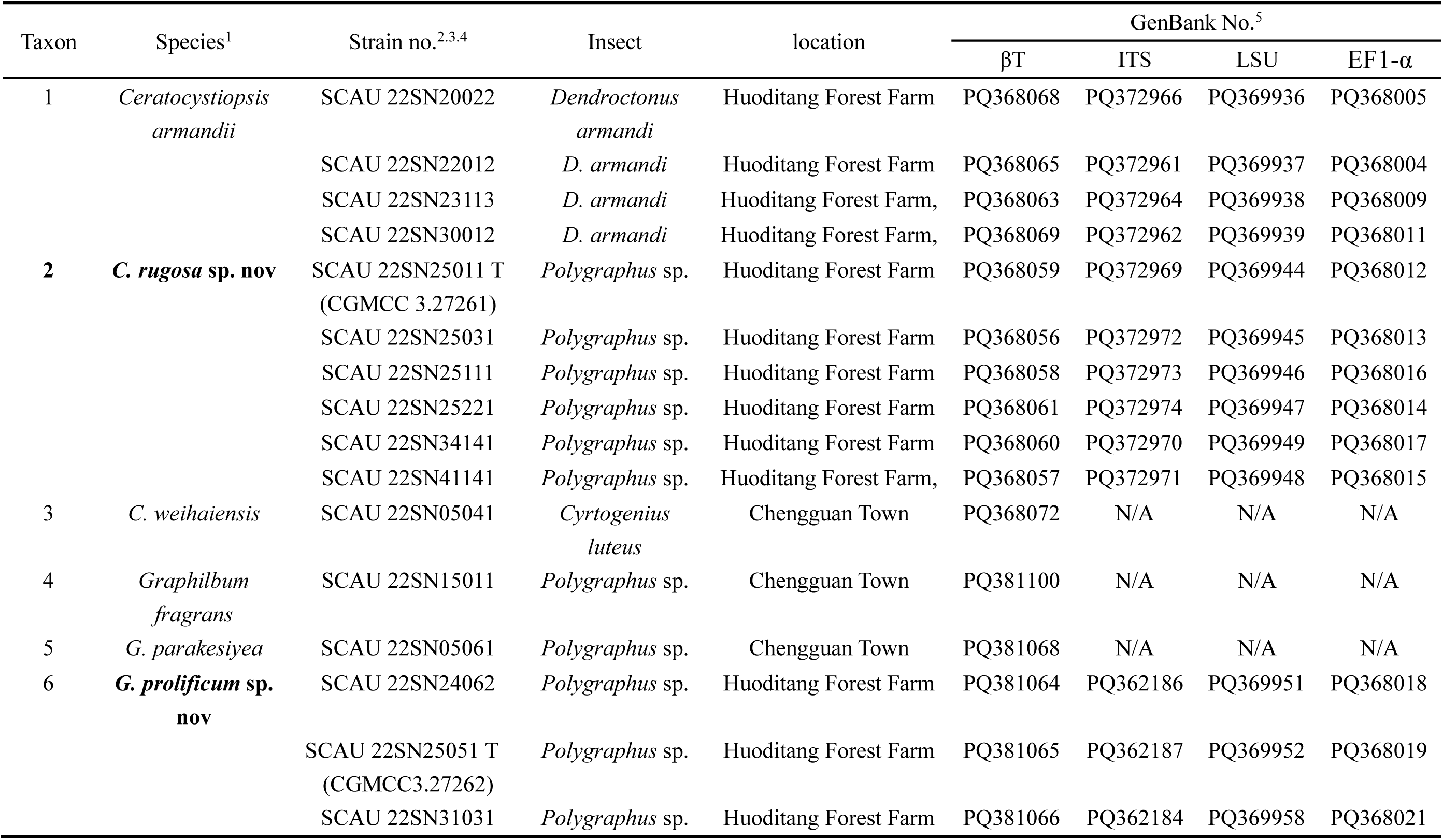
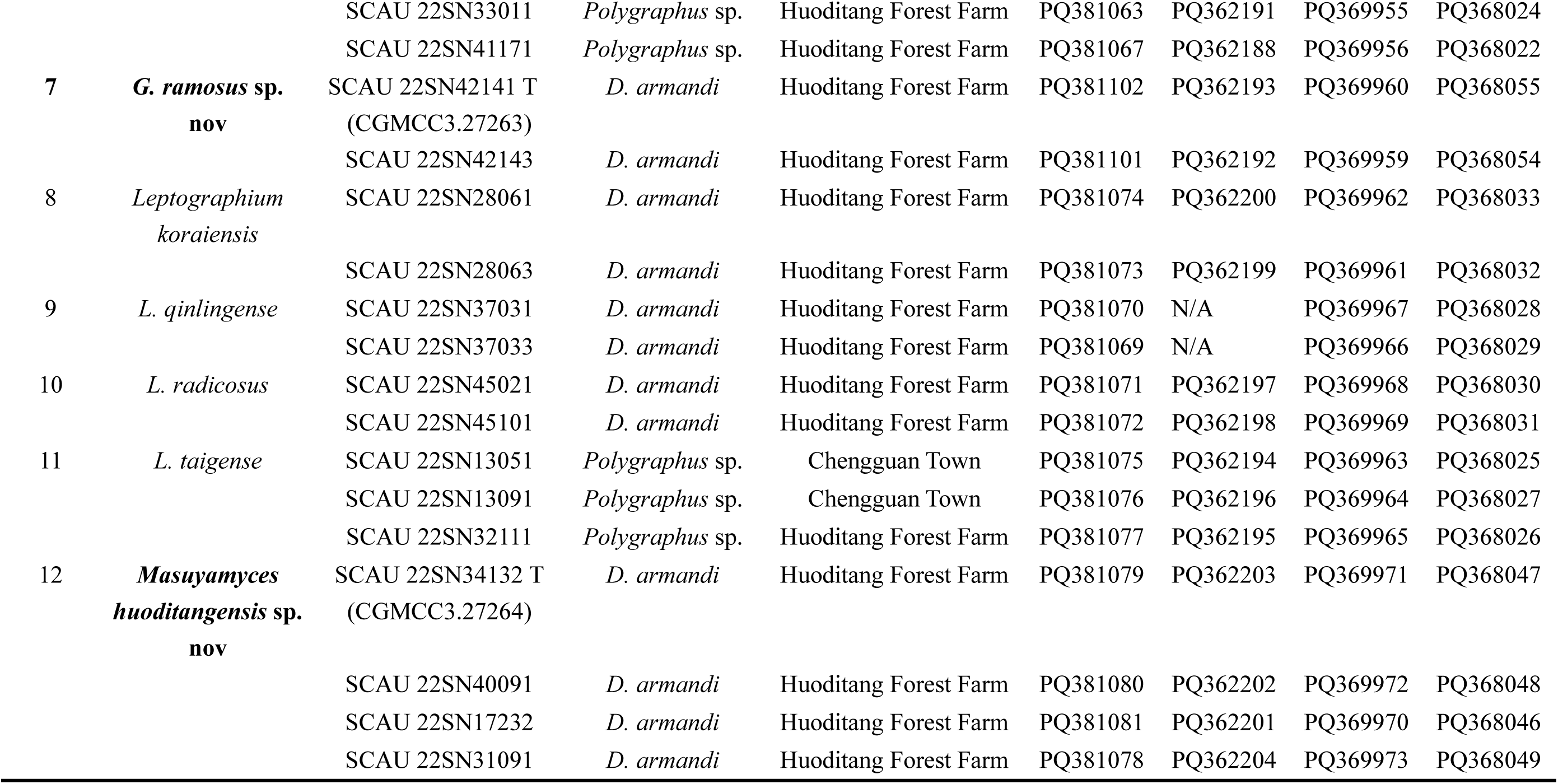
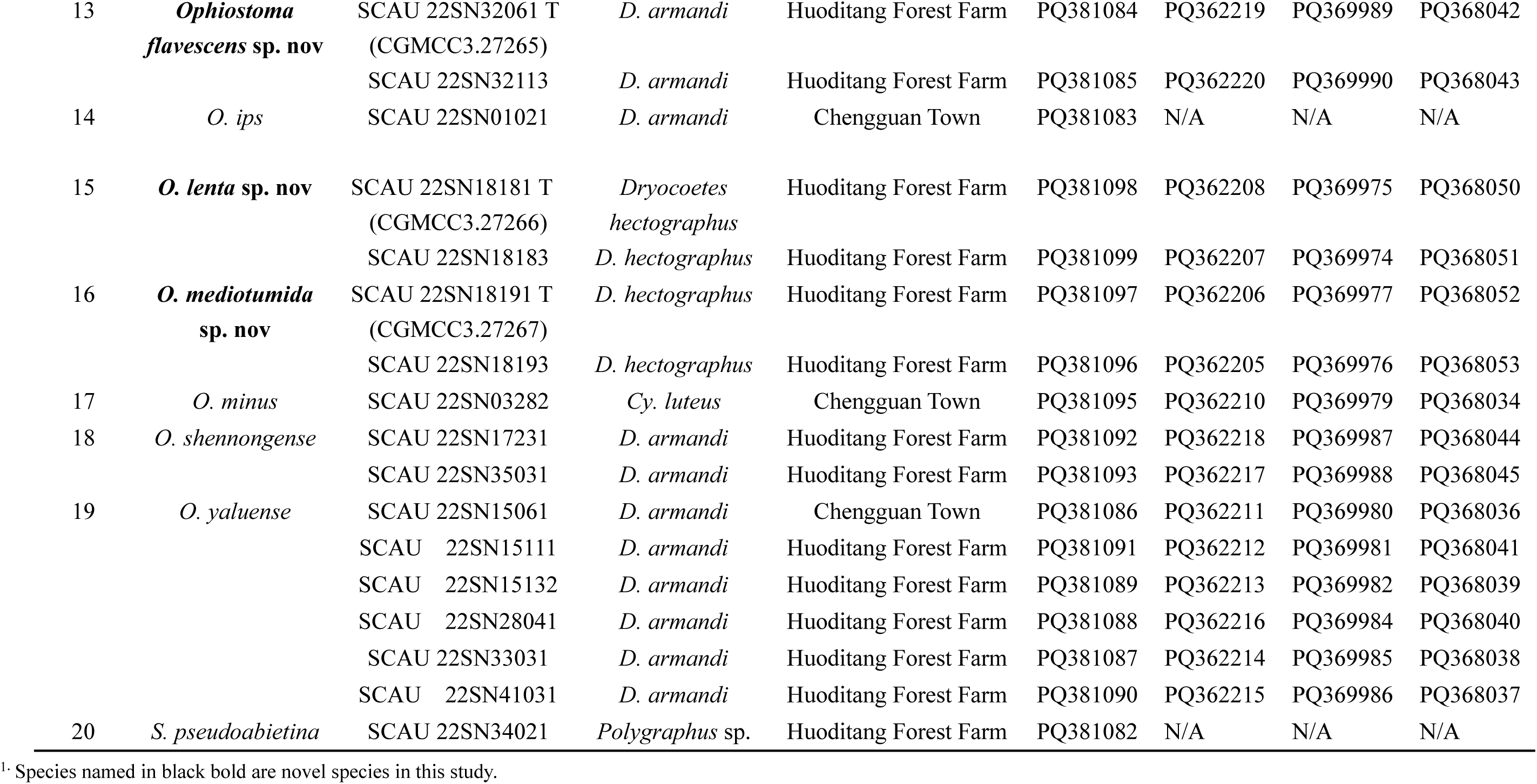
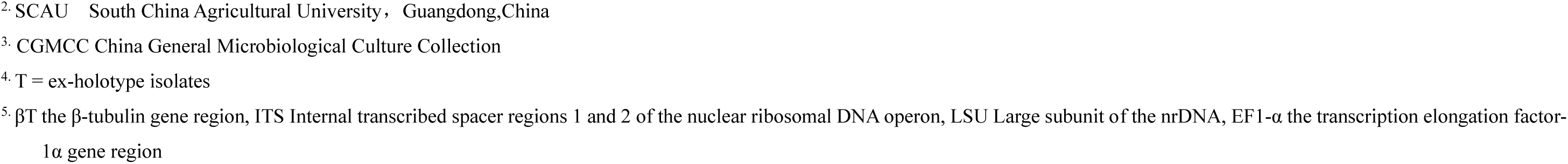
Representative isolates of ophiostomatoid fungi associated with bark beetles Infesting P. armandii obtained in this study.

### DNA extraction and PCR amplification

Before DNA extraction, the strain was grown on 2 % MEA at 25 ° C in the dark for 1-2 weeks. The viable mycelium at the tip of the pure strain was scraped to an appropriate size of 1.5 ml sterile centrifuge tubes. According to the instructions, DNA was extracted using PrepMan TM Ultra Sample Preparation Reagent (by Thermo Fisher Scientific). A total of 4 DNA regions were amplified for sequencing and phylogenetic analysis. BT2a and BT2b primer pairs amplified the β-tubulin (TUB2) gene; ITS5 and ITS4 primer pairs amplified the internal transcribed spacer (ITS) regions; the ribosomal large subunit (LSU) region was amplified by LROR and LR5 primer pairs; and EF2F and EF2R primer pairs amplified the elongation factor 1-α (EF1-α) gene.

PCR detection was performed in 25 μL (12.5 μL 2 × Taq Master Mix, 0.5 μL forward and reverse primers, 10.5 μL PCR grade deionized water, and one μL DNA stock solution ). The PCR amplification conditions were pre-denaturation at 95 °C for 3 minutes, then denaturation at 95 °C for 30 seconds, annealing at 52 °C - 58 °C for 30 seconds, extension at 72 °C for 1 minute, a total of 35 cycles, and finally extension at 72 °C for 10 minutes, stored at 12 °C.

All PCR products were sent to Sangon Biotechnology Co., Ltd. in Guangzhou, Guangdong Province, China, for sequencing, and the sequences were downloaded from Sangon’s website (https://store.sangon.com) after the sequencing was completed.

### Phylogenetic Analyses

The downloaded sequences were assembled by Geneious v. 7.1.4 (Biomatters, Auckland, New Zealand). The sequences obtained from the four DNA fragments were preliminarily identified by BLAST search in the NCBI GenBank database. The representative sequences with the highest similarity matching and the model strain sequences of similar species were downloaded from the NCBI GenBank database. The GenBank numbers of these sequences are shown on the corresponding phylogenetic trees. The dataset was assembled in MEGA v. 7.0 (Kumar et al., 2016) and compared using the online tool MAFFT v. 7 (Katoh et al., 2019). Phylogenetic analysis was performed using maximum likelihood (ML), maximum parsimony (MP), and Bayesian inference (BI).

The best ML, MP, and BI alternative models were selected in JModelTest 2.1.10 (Darriba et al., 2012). MEGA-X (Kumar et al., 2018) is used for ML analysis of the nearest neighbor exchange (NNI) branch exchange options. One thousand guided replications determine the confidence interval of the node. MP analysis uses PAUP * Version 4.0b10 (Swofford, 2002) and performs 1000 times guided replication to determine the credibility of the branch node. BI analysis was performed using MrBayes V.3.2.7 (Ronquist et al., 2003), starting from a random starting tree, running a total of four Markov chain Monte Carlo (MCMC) chains, and running 5 million random generations at the same time to calculate the posterior probability, using the best alternative model identified in jModelTest 2.1.10. Sampling once every 100 generations, 50,000 trees were obtained. In addition, the first 25 % of the sampled trees were discarded during the combustion process, and the retained tree calculated the posterior probability. The phylogenetic tree was edited in FigTree V.1.4.2.

### Morphological and physiological characteristics

Agar blocks with a side length of 5 mm were selected from the colonies with vigorous growth activity, placed in the center of a culture dish with a diameter of 90 mm containing 2 % MEA, and cultured in a dark environment at 25 °C. Each strain replicated five plates to study the growth rate of the colony. The two orthogonal diameters of the strains were recorded on the 7th and 14th day of colony growth. According to Rayner’s color card (1970), the colony color is described. All data related to the type specimen are stored in MycoBank.

For the microstructure of the new species, the strain was first inoculated on a pine strip plate and cultured at 25 °C in a dark environment. The morphology and growth status of the culture were observed daily, and then the pine strip plate cultured for 2-3 weeks was made into slides to observe the sexual or asexual structure. Zeiss Axio Imager Z2 (Carl Zeiss, Germany) was used to measure and photograph the microstructure of ophiostomatoid fungi. The structure of each model strain, such as conidia and conidiophores, was measured 30 times.

## RESULTS

### Collection of samples and isolation of fungi

This study isolated 290 strains of fungi, including 225 strains of Crustacea fungi (77 strains from Chengguan Town, Shaanxi Province, and 148 strains from Huoditang Forest Farm, Shaanxi Province). The growth rate and macroscopic and microscopic morphological characteristics were used for preliminary identification. The βT sequences of all strains were used for standard nucleotide BLAST search in GenBank to facilitate preliminary classification and affinity search. Then, 50 representative strains were selected for in-depth morphological study and multi-locus phylogenetic analysis (Table 1).

### Phylogenetic analysis

Similar topological structures are generated using ML, MP, and BI phylogenetic methods and the statistical support of each sequence data set changes slightly. The phylogenetic tree of each dataset is obtained by ML analysis, and the branch support is obtained by ML, MP, and BI analyses. The best evolutionary models selected by jModelTest v. 2.1.10 were GTR + G (for the ITS dataset of *Graphilbum*), GTR + I + G[for the EF1-α and LSU dataset of *Graphilbum*, the EF1-α dataset of *O. clavatum* complex, the combined datasets (βT + ITS +LSU+ EF1-α)of *Ceratocystiopsis*, the combined datasets (βT + ITS) of *Graphilbum*, the combined datasets (βT + ITS + EF1- α) of *Leptographium*, the combined datasets (ITS + LSU + EF1-α)of *Masuyamyces*, the combined datasets (βT+ ITS + EF1-α)of *O. clavatum* complex], TrN + I + G (for the βT dataset of ophiostomatoid, the ITS and EF1-α dataset of *Ceratocystiopsis*, the EF1-α dataset of *Leptographium* and *Masuyamyces*, the LSU dataset of *Leptographium* and *Masuyamyces*), TrN + G (for the ITS dataset of *Leptographium*, *Masuyamyces* and *O. clavatum* complex, the LSU dataset of *Ceratocystiopsis*), TrN+I (for the LSU dataset of *O. clavatum* complex).

In the ophiostomatoid fungi βT sequence data set, the βT fragment is 453 bp long, including the gap. Phylogenetic analysis showed that the strains produced in this study belonged to 20 terminal clades or phylogenetic species, and three phylogenetic species were nested in *Ceratocystiopsis* (taxa 1-3). Four phylogenetic species nested in *Graphilbum* (taxa 4-7), three phylogenetic species nested in *Leptographium* ( taxa 8-10 ), one phylogenetic species nested in *Masuyamyces* (taxa 12), five phylogenetic species nested in *Ophiostoma* (taxa 13-14,17-19), and the last phylogenetic species nested in *Sporothrix* (taxa 20). Furthermore, the three taxa are distributed outside the currently recognized species complex but belong to previously shown“Group A”( taxa 15-16 ) (Runlei et al., 2017) and “Group C” ( taxa 11 ) ( Chang et al., 2019 ).

#### Ceratocystiopsis

The ITS, EF1-α, LSU dataset, and combined datasets ( βT + ITS + LSU + EF1-α ) of *Ceratocystiopsis* comprised 558,655,538 and 2393 characters, respectively, including gaps. According to the phylogenetic analysis of the βT gene region ( Fig. 2 ), the four representative strains of taxon one and one representative strain of taxon 3 were identical to the sequences of *C. armandii* and *C. weihaiensis*, respectively, and clustered into the same clade ( Fig. 2 ), so taxon two and taxon three were defined as previously known species. The phylogenetic analysis of LSU ( Fig. 4 ) showed that the six representative strains of taxon two were clustered in the same subbranch with *C. hongchibaensis* and *C. debeeria*, but LSU could not distinguish closely related species in all cases. Therefore, combined with the phylogenetic analysis of βT, ITS, EF1-α, and combined datasets ( βT + ITS + LSU + EF1-α ) ( Fig. 2-3, 5-6 ), taxon 2 formed a suitable branch. It is most closely related to *C. hongchibaensis* and *C. debeeria*.

**Fig. 2.**
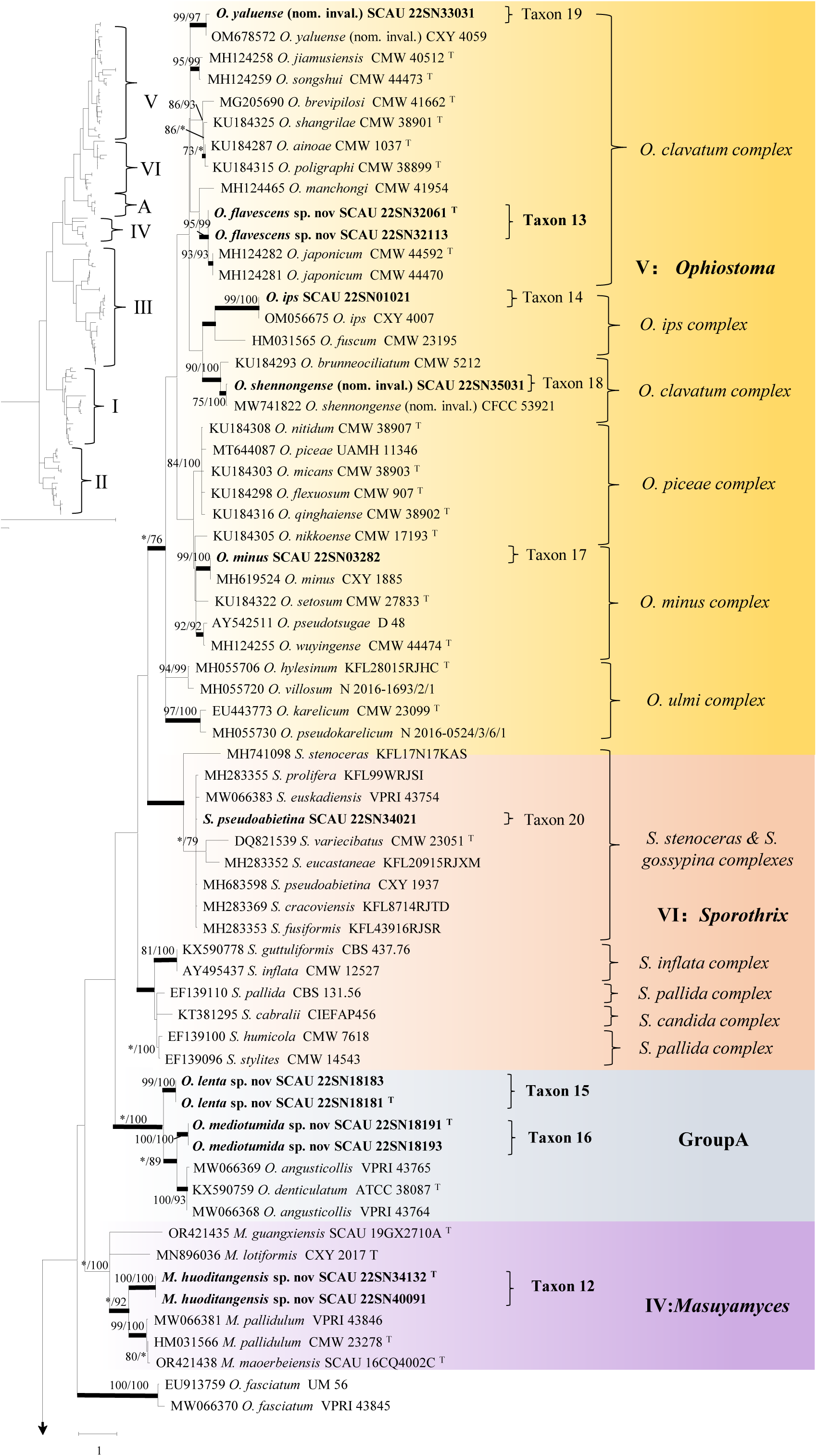
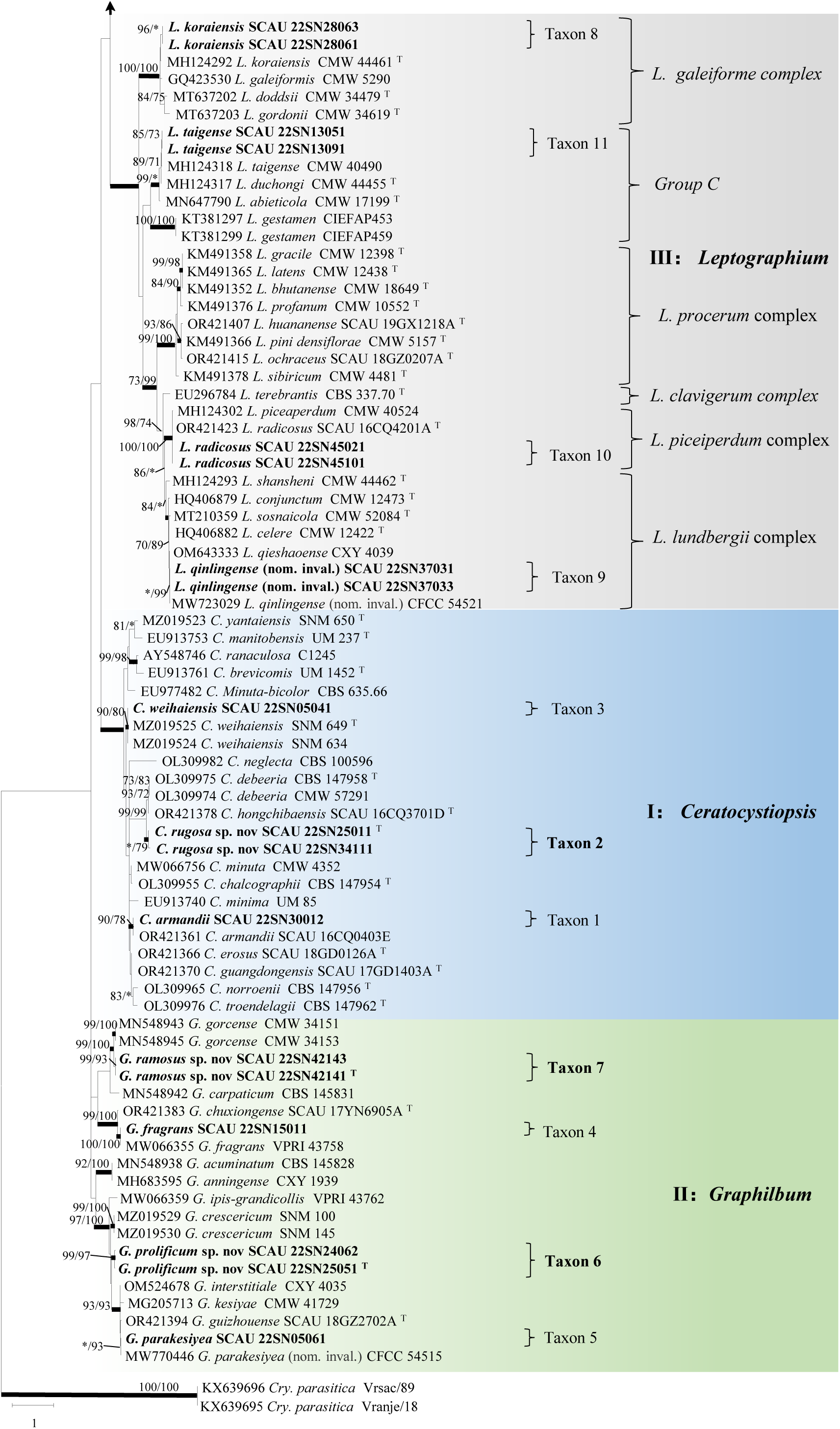
ML tree of ophiostomatoid fungi from the BT sequence data. Sequences generated from this study are printed in bold. Bold branches indicate posterior probability values ≥0.95. Bootstrap values of ML/MP ≥ 70% are recorded at the nodes. T = ex-type isolates.

**Fig. 3.**
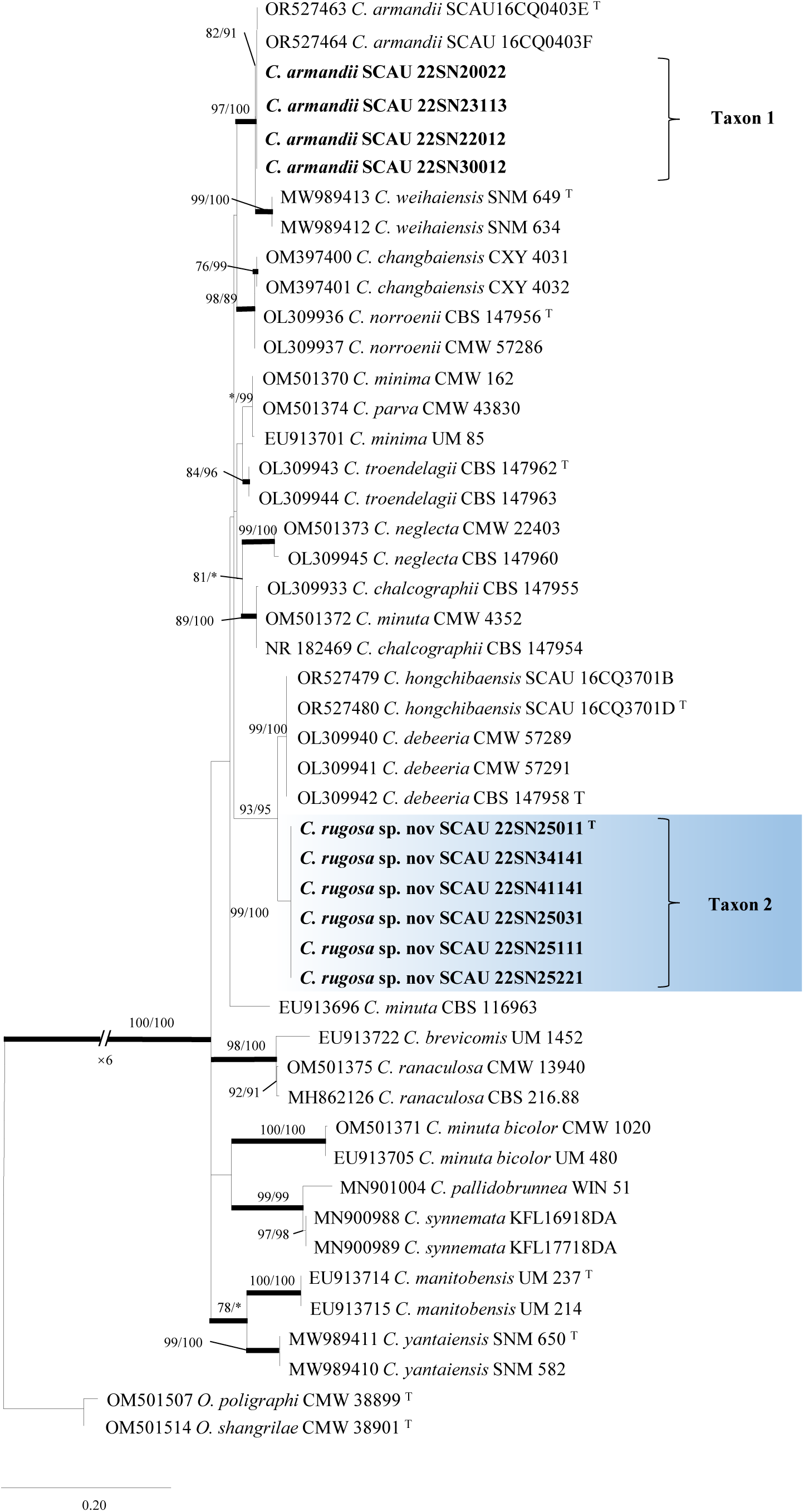
ML tree of *Ceratocystiopsis* generated from the ITS sequence data. Sequences generated from this study are printed in bold. Bold branches indicate posterior probability values ≥0.95. Bootstrap values of ML/MP ≥ 70% are recorded at the nodes. T = ex-type isolates.

**Fig. 4.**
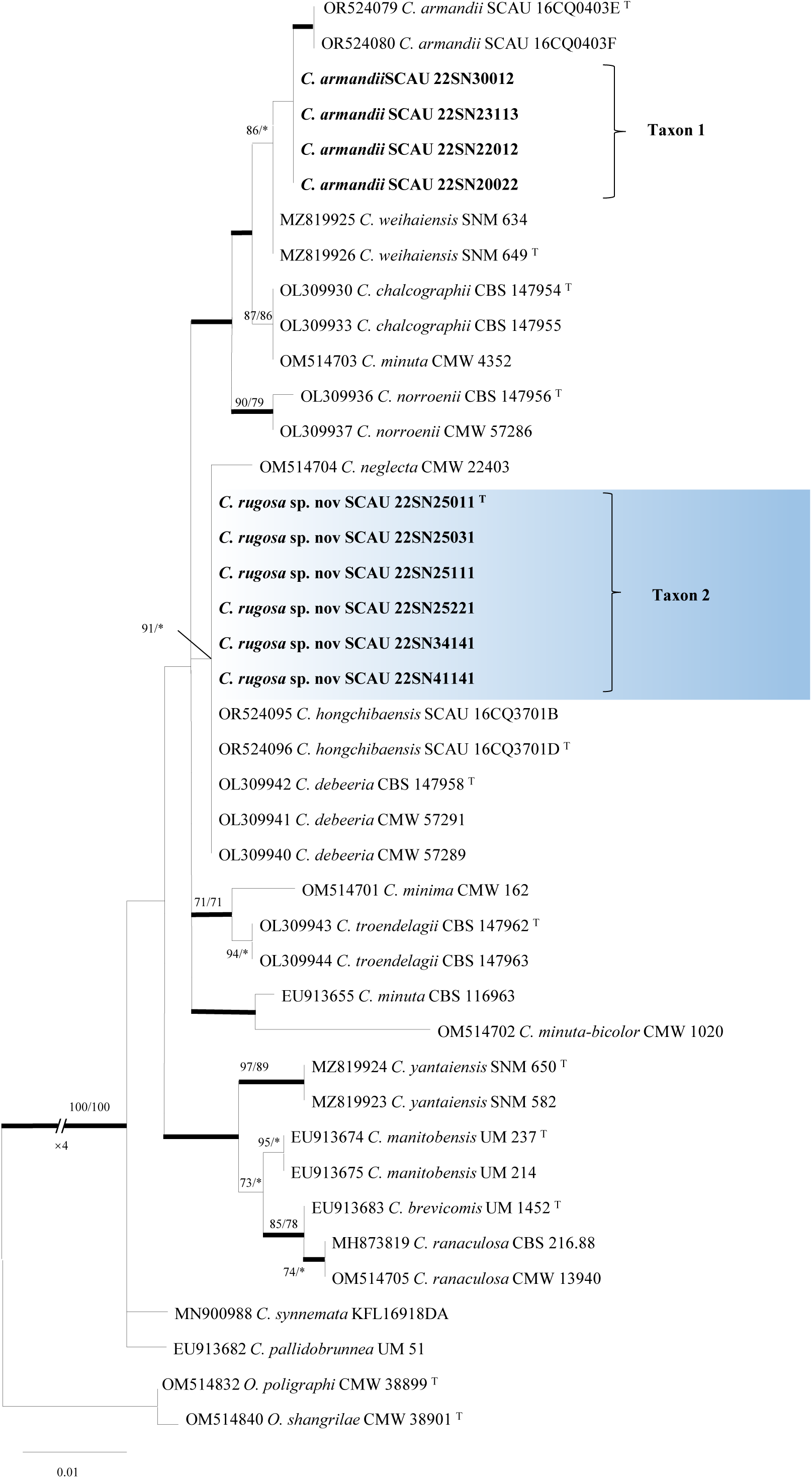
ML tree of *Ceratocystiopsis* generated from the LSU sequence data. Sequences generated from this study are printed in bold. Bold branches indicate posterior probability values ≥0.95. Bootstrap values of ML/MP ≥ 70% are recorded at the nodes. T = ex-type isolates.

**Fig. 5.**
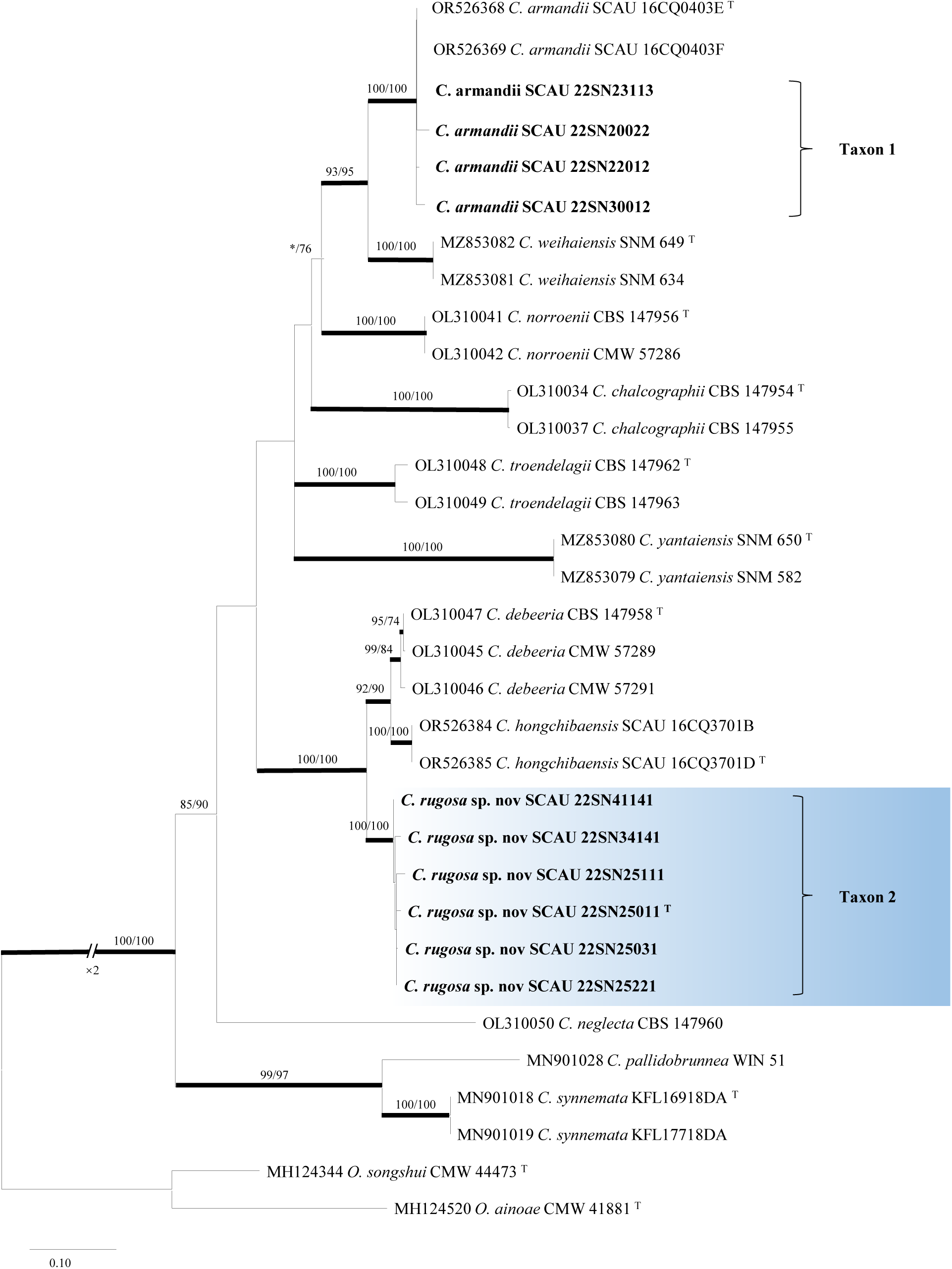
ML tree of *Ceratocystiopsis* generated from the EF1-α sequence data. Sequences generated from this study are printed in bold. Bold branches indicate posterior probability values ≥0.95. Bootstrap values of ML/MP ≥ 70% are recorded at the nodes. T = ex-type isolates.

**Fig. 6.**
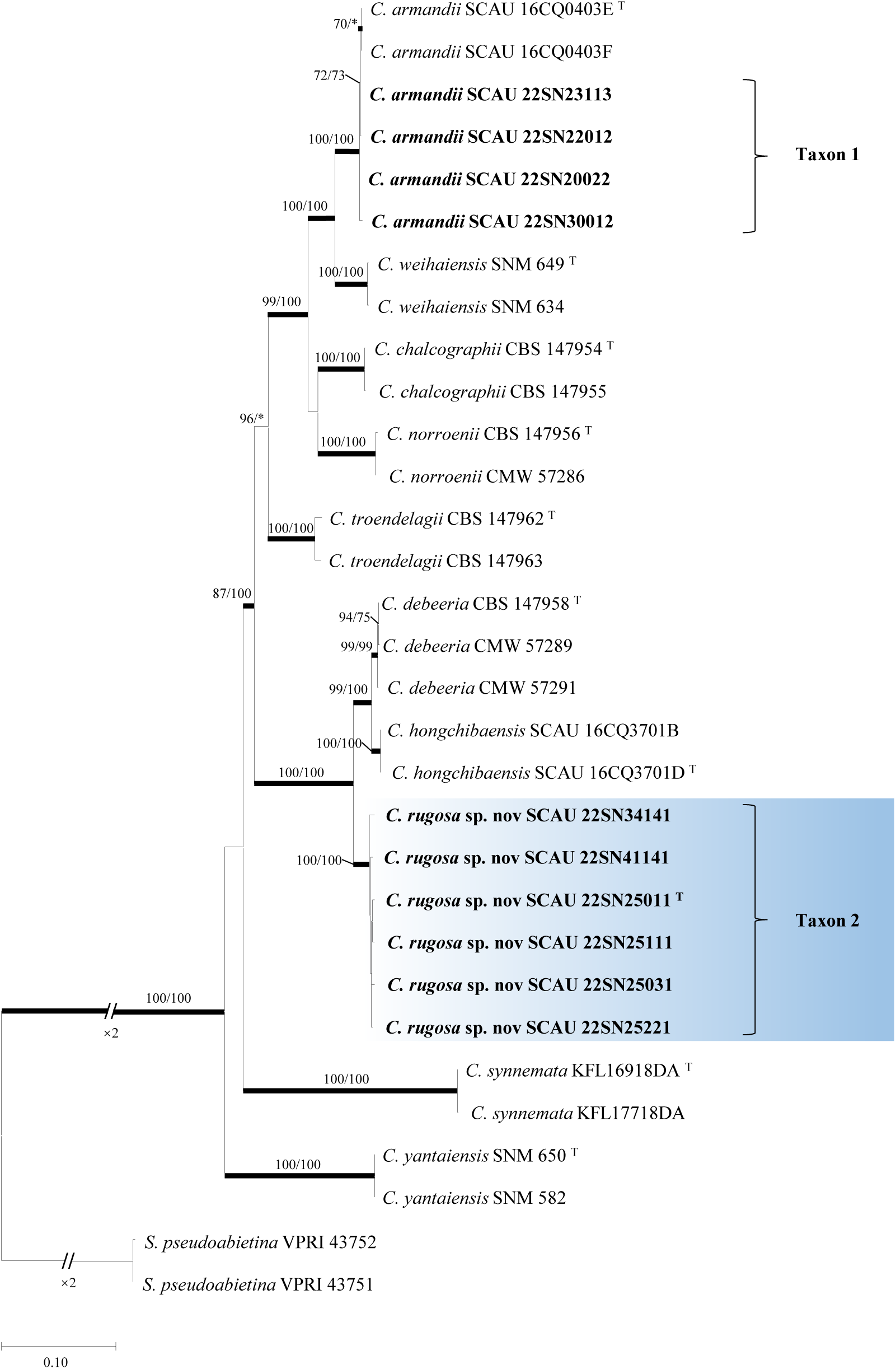
ML tree of *Ceratocystiopsis* generated from the combined (βT+ ITS +LSU+ EF1-α) sequence data. Sequences generated from this study are printed in bold. Bold branches indicate posterior probability values ≥0.95. Bootstrap values of ML/MP ≥ 70% are recorded at the nodes. T = ex-type isolates.

#### Graphilbum

The ITS, LSU, EF1-α datasets and the combined dataset ( βT + ITS ) of the genus *Graphilbum* are composed of 522, 559, 512 and 1209 characters, respectively, including gaps. The five representative strains of taxon 6 and two representative strains of taxon 7 formed a branch with high node support in the phylogenetic tree of βT, ITS, LSU EF1-α and combined dataset ( βT + ITS ) ( Fig. 2, 7-10 ). Taxon 6 was closely related to *G. sexdentatum* and *G. crescericum*, and taxon 7 was closely related to *G. gorcense*. One representative strain of taxon 4 and one representative strain of taxon 5 were the same as the sequences of *G. fragrans* and *G. parakesiyea* and clustered in the same branch in the phylogenetic analysis of βT ( Fig. 2 ), so the taxon was determined to be a previously discovered known species.

**Fig. 7.**
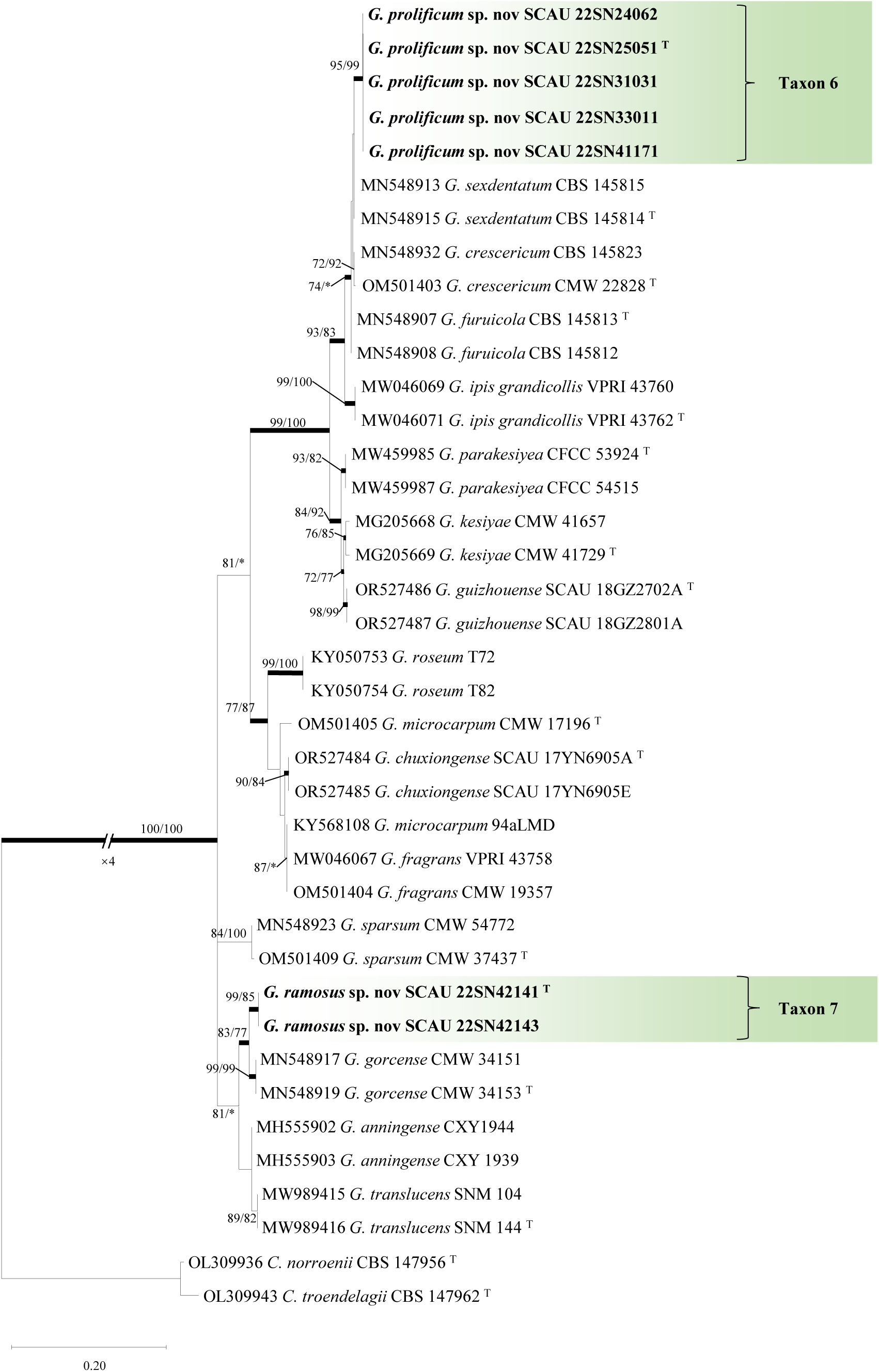
ML tree of *Graphilbum* generated from the ITS sequence data. Sequences generated from this study are printed in bold. Bold branches indicate posterior probability values ≥0.95. Bootstrap values of ML/MP ≥ 70% are recorded at the nodes. T = ex-type isolates.

**Fig. 8.**
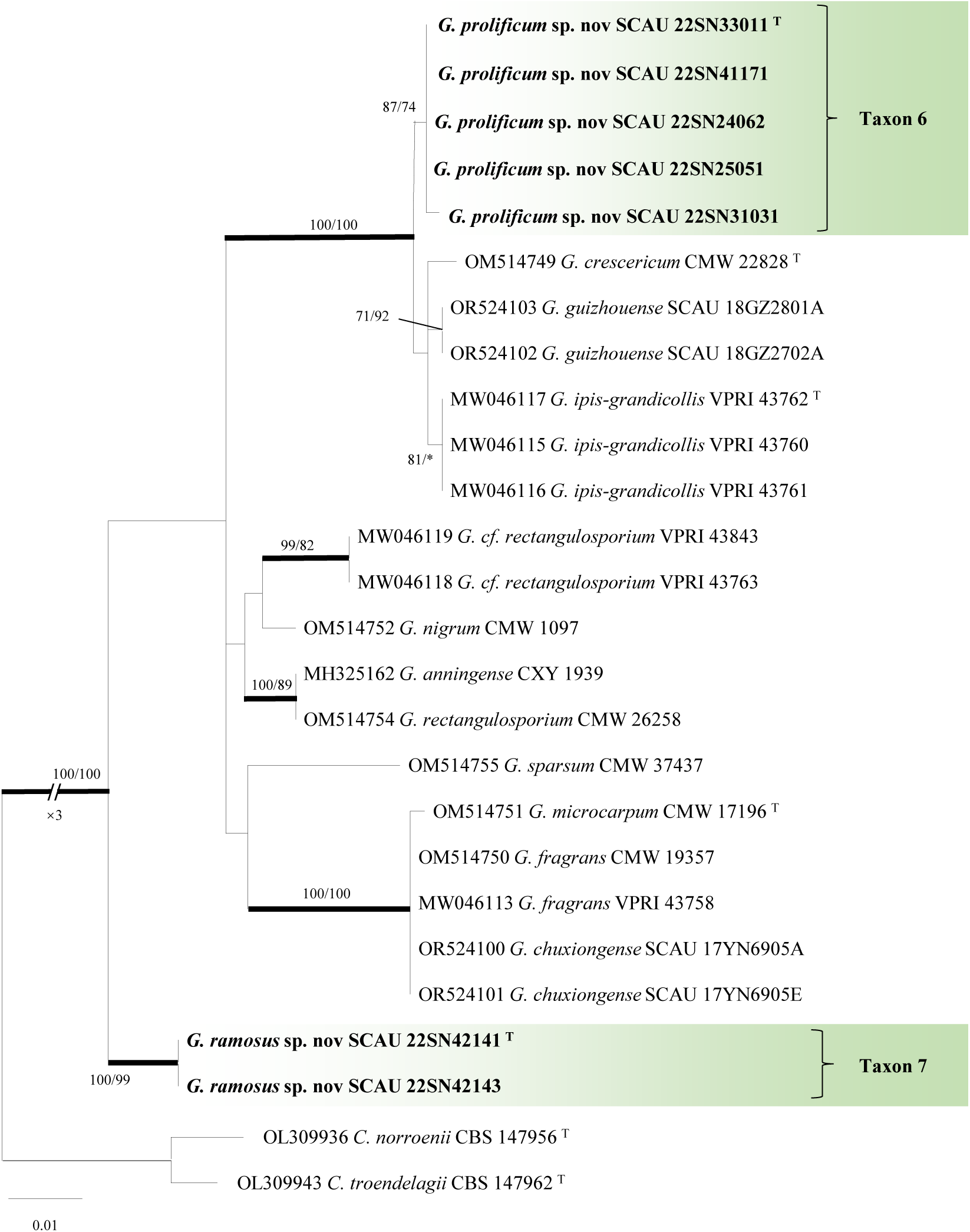
ML tree of *Graphilbum* generated from the LSU sequence data. Sequences generated from this study are printed in bold. Bold branches indicate posterior probability values ≥0.95. Bootstrap values of ML/MP ≥ 70% are recorded at the nodes. T = ex-type isolates.

**Fig. 9.**
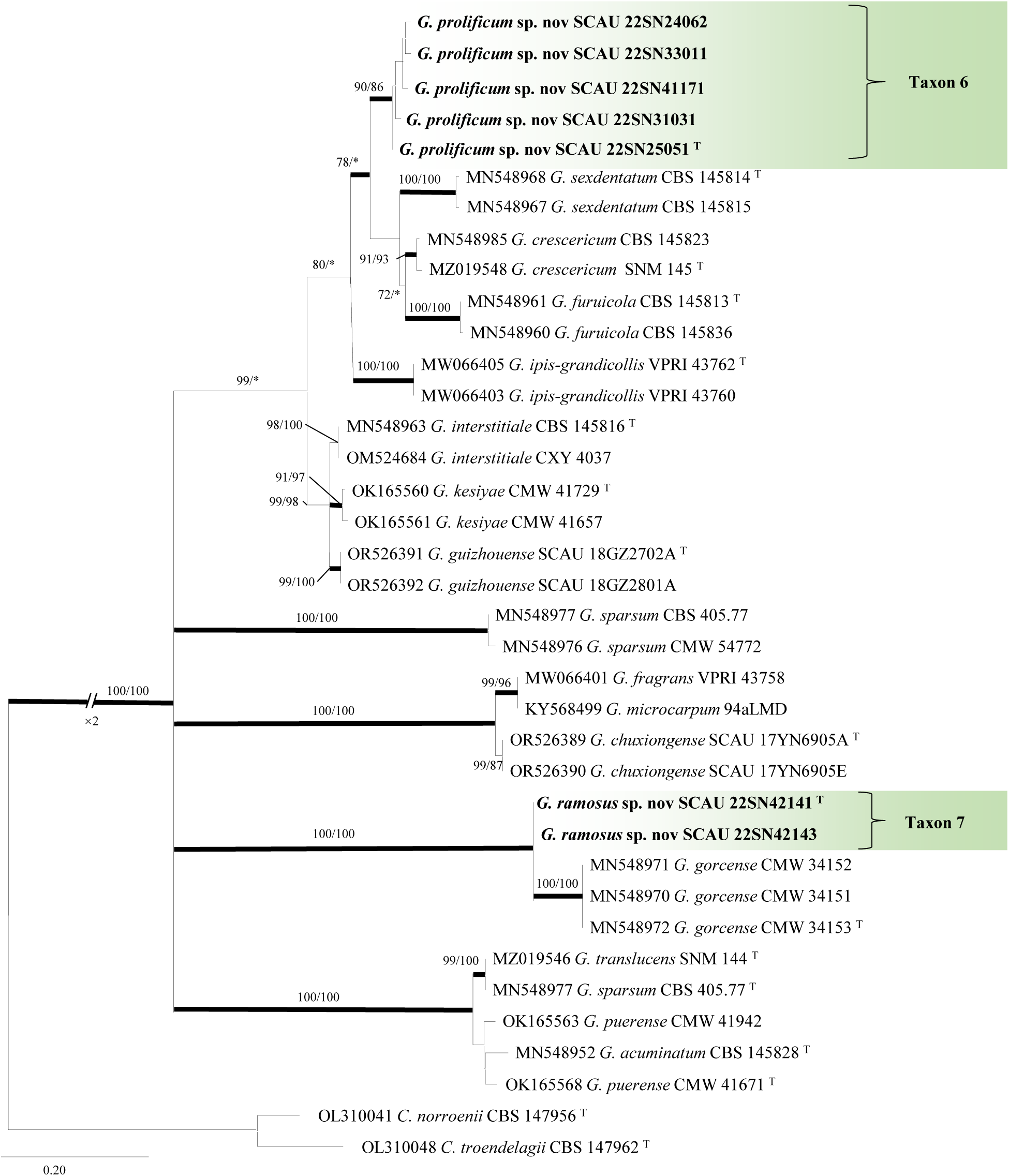
ML tree of *Graphilbum* generated from the EF1-α sequence data. Sequences generated from this study are printed in bold. Bold branches indicate posterior probability values ≥0.95. Bootstrap values of ML/MP ≥ 70% are recorded at the nodes. T = ex-type isolates.

**Fig. 10.**
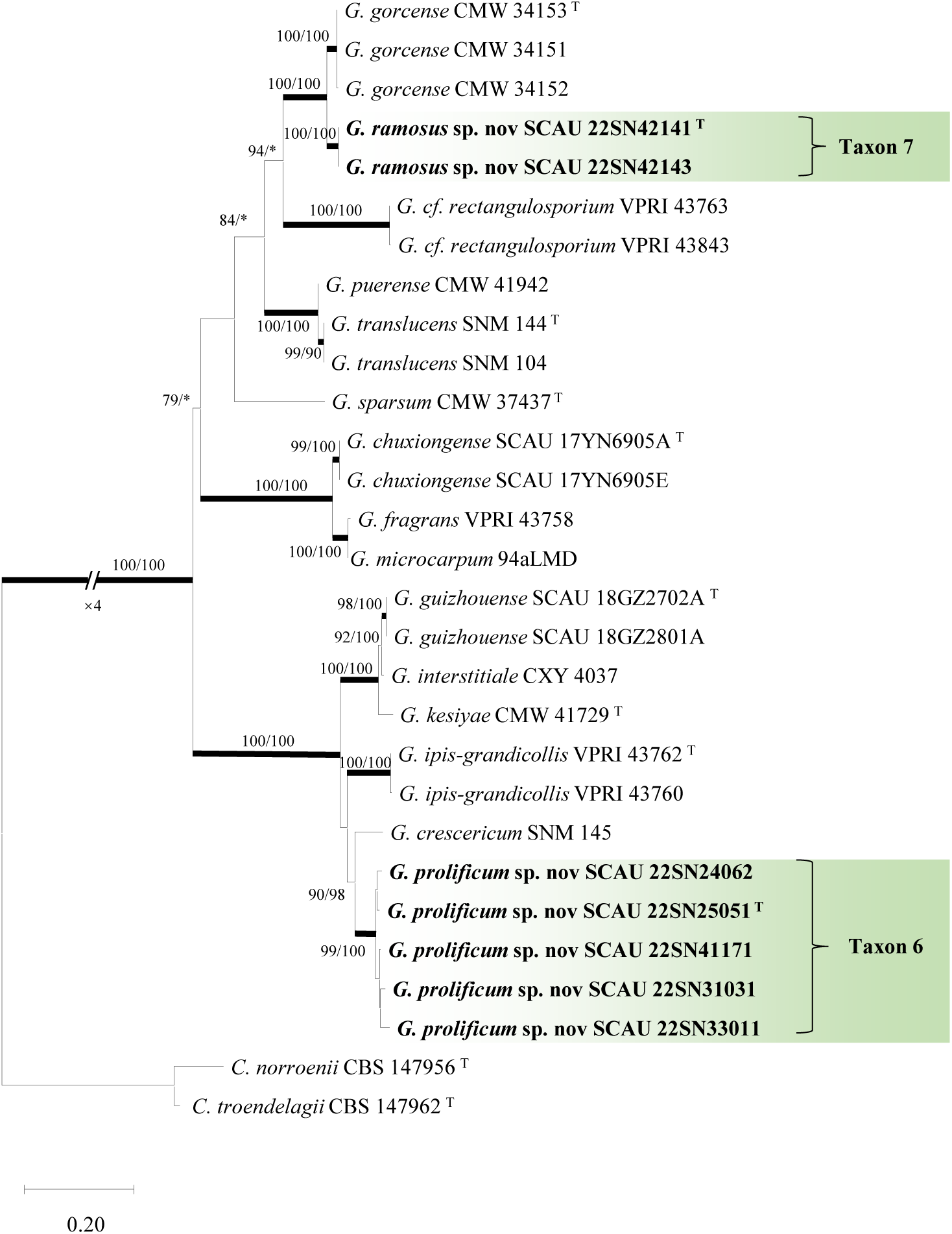
ML tree of *Graphilbum* generated from the combined (βT+ ITS)sequence data. Sequences generated from this study are printed in bold. Bold branches indicate posterior probability values ≥0.95. Bootstrap values of ML/MP ≥ 70% are recorded at the nodes. T = ex-type isolates.

#### Leptographium and “Group C”

The ITS, LSU, EF1-α and combined dataset ( βT + ITS + EF1-α ) of *Leptographium* and “GroupC”were composed of 537, 703, 666 and 1412 characters, respectively, including gaps. The two representative strains of taxon 8 formed an independent branch in the phylogenetic analysis of βT, LSU, EF1-α and the combined dataset ( βT + ITS + EF1-α ) ( Fig. 2, 12-14 ), while the inference based on ITS phylogenetic analysis ( Fig. 11 ) showed that the two strains were clustered with *L. koraiensis*. According to the data sets of different fragments, the support of the independent branch nodes formed by taxon 8 is relatively low, and the base sequence is basically consistent with the *L. koraiensis* model strain, so it is judged to be a known species. The two representative strains of taxon 9 belong to the *L. lundbergii* complex in the βT phylogenetic tree ( Fig. 2 ). Combined with the phylogenetic analysis of LSU and EF1-α ( Fig. 12-13 ), taxon 9 and *L. qinlingense* are clustered in the same branch. Due to *L. qinlingense* lacks the type specimen, it is considered an invalid name. The two representative strains of taxon 10 belong to the *L. piceiperdum* complex in the BT phylogenetic analysis ( Fig. 2 ). Combined with the phylogenetic analysis of ITS, LSU, EF1-α and combined dataset ( βT + ITS + EF1-α ) ( Fig. 11-14 ), taxon 10 and *L. radicosus* are clustered in the same branch and have high support, so it is determined that taxon 10 is a previously discovered known species. Although the three representative strains of taxon 11 formed independent branches in the phylogenetic analysis of βT and EF1-α ( Fig. 2, 13 ), combined with the phylogenetic development of ITS and LSU ( Figs. 11-12 ), taxon 11 and *L. taigense* were clustered in the same branch with high support, so it was determined that taxon 11 was a known species in the previously discovered “ Group C ” ( Chang et al. 2019 ).

**Fig. 11.**
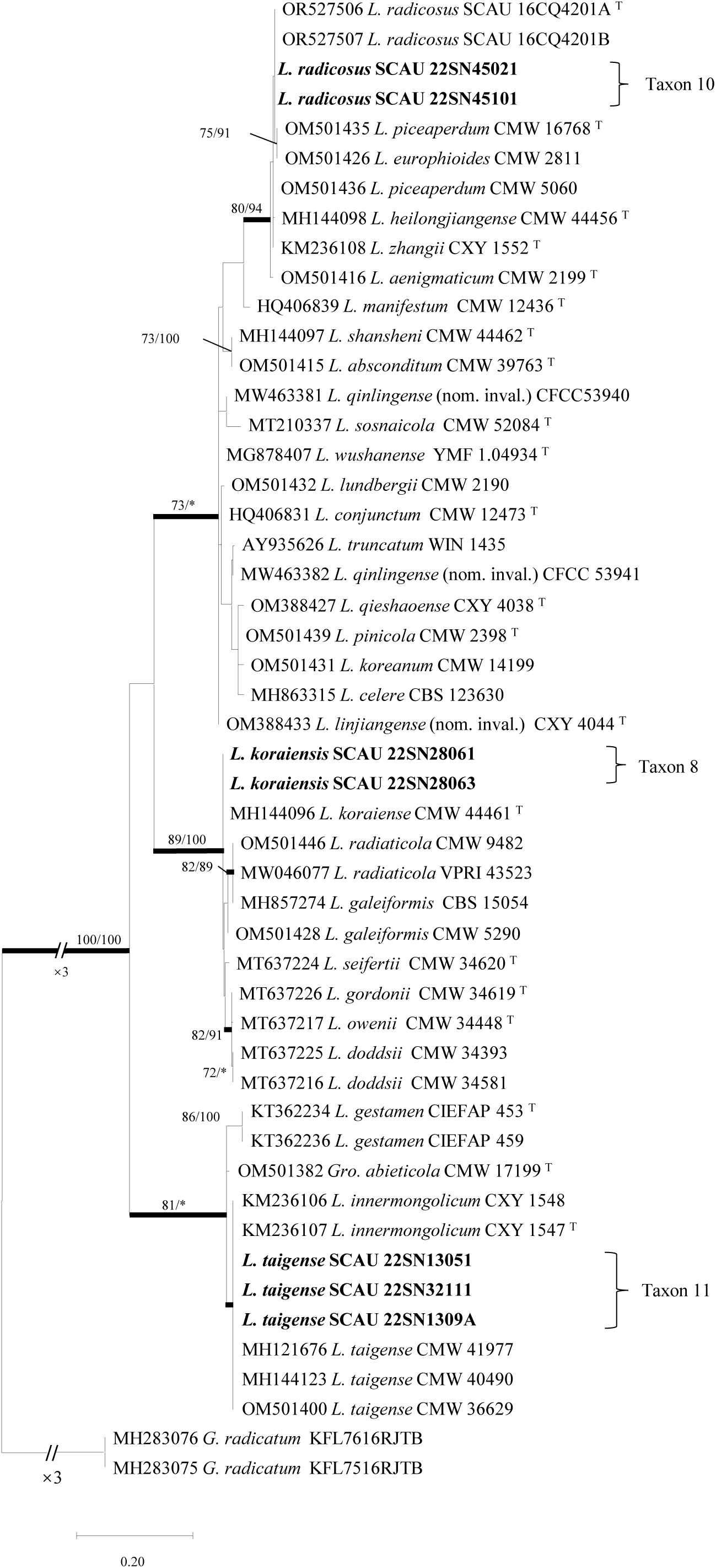
ML tree of *Leptographium* and“Group C”generated from the ITS sequence data. Sequences generated from this study are printed in bold. Bold branches indicate posterior probability values ≥0.95. Bootstrap values of ML/MP ≥ 70% are recorded at the nodes. T = ex-type isolates.

**Fig. 12.**
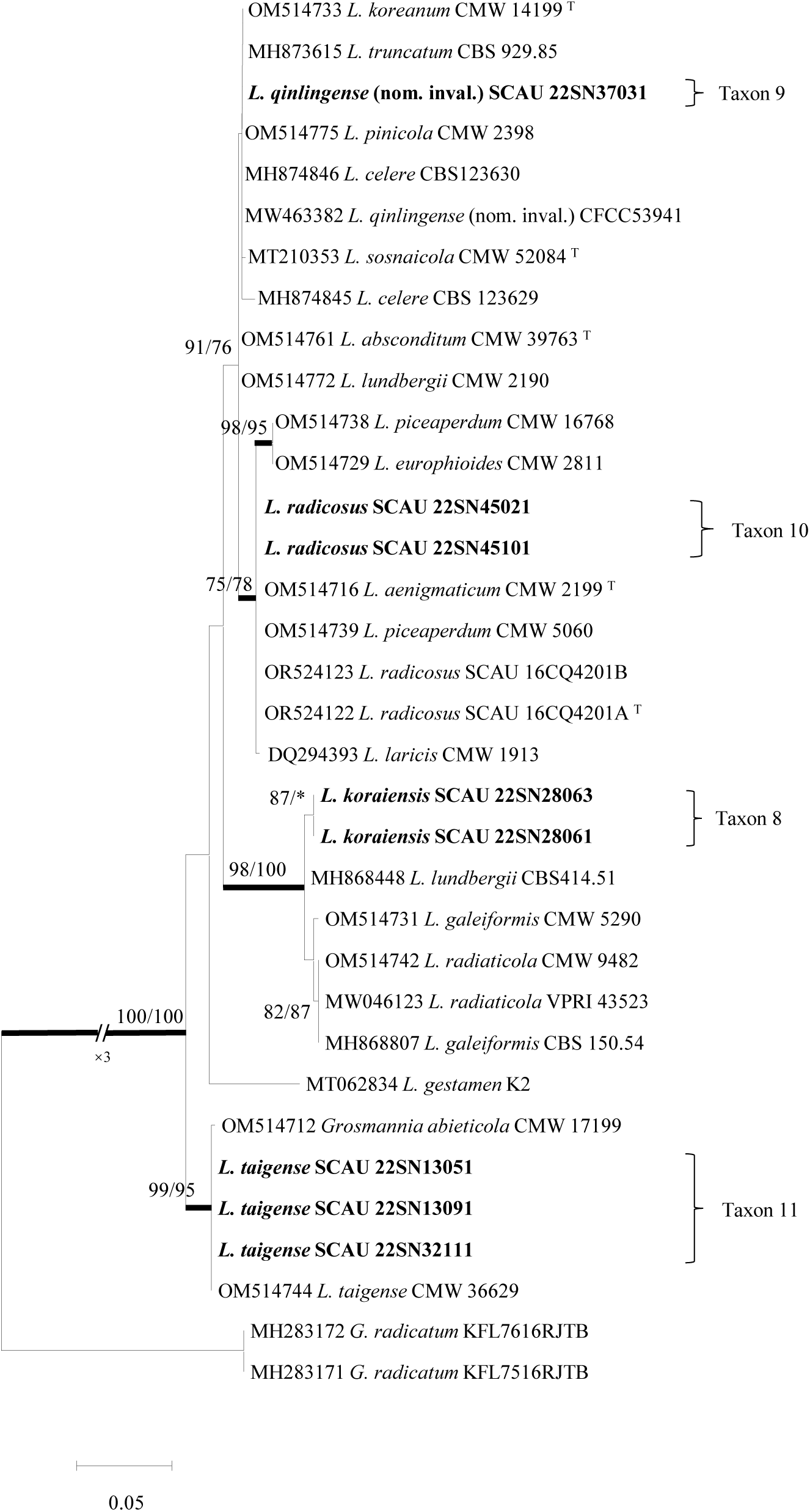
ML tree of *Leptographium* and“Group C”generated from the LSU sequence data. Sequences generated from this study are printed in bold. Bold branches indicate posterior probability values ≥0.95. Bootstrap values of ML/MP ≥ 70% are recorded at the nodes. T = ex-type isolates.

**Fig. 13.**
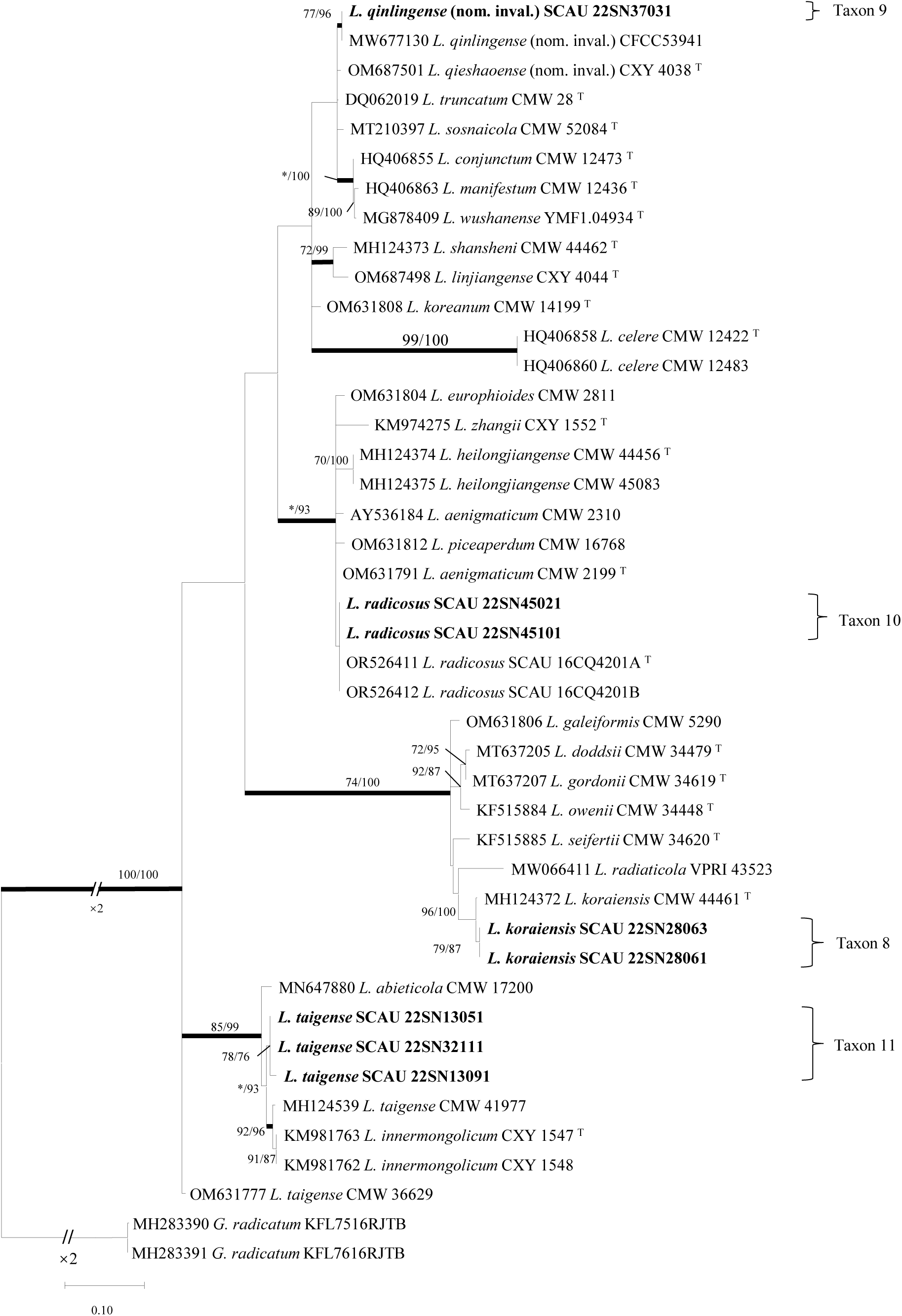
ML tree of *Leptographium* and“Group C”generated from the EF1-α sequence data. Sequences generated from this study are printed in bold. Bold branches indicate posterior probability values ≥0.95. Bootstrap values of ML/MP ≥ 70% are recorded at the nodes. T = ex-type isolates.

**Fig. 14.**
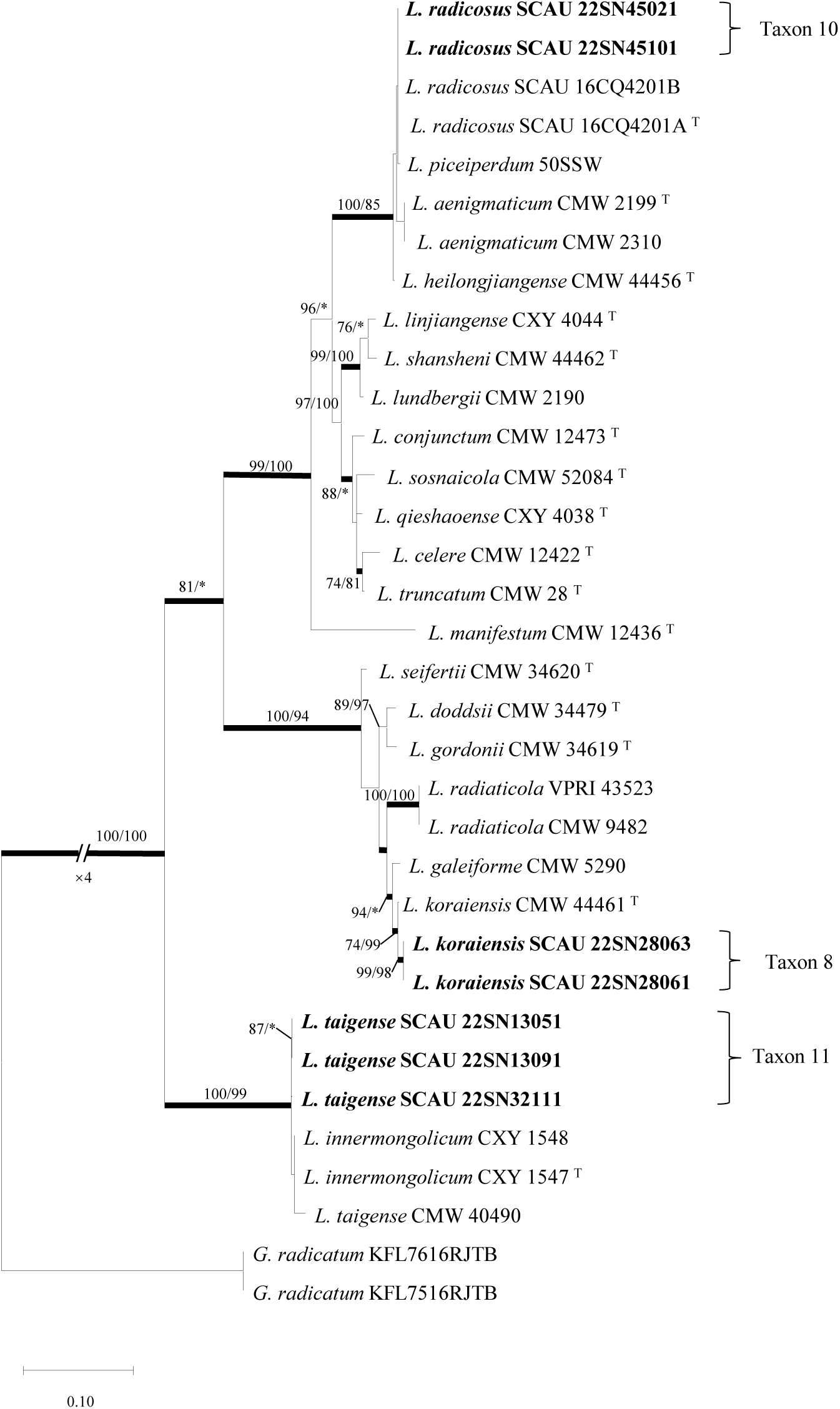
ML tree of *Leptographium* and“Group C”generated from the combined (βT + ITS + EF1-α) sequence data. Sequences generated from this study are printed in bold. Bold branches indicate posterior probability values ≥0.95. Bootstrap values of ML/MP ≥ 70% are recorded at the nodes. T = ex-type isolates.

#### *Masuyamyces* and “Group A”

The ITS, LSU, EF1-α datasets and the combined dataset ( ITS + LSU + EF1-α ) of Masuyamyces are composed of 535, 768, 736 and 1715 characters, respectively, including gaps. The four representative strains of taxon 12 formed a well-supported branch in the phylogenetic tree of βT, ITS, LSU, EF1-α and the combined dataset ( ITS + LSU + EF1-α ) ( Fig. 2, 15-18 ), which is closely related to *M. pallidulum* but different, so it is considered to be a unique species.

**Fig. 15.**
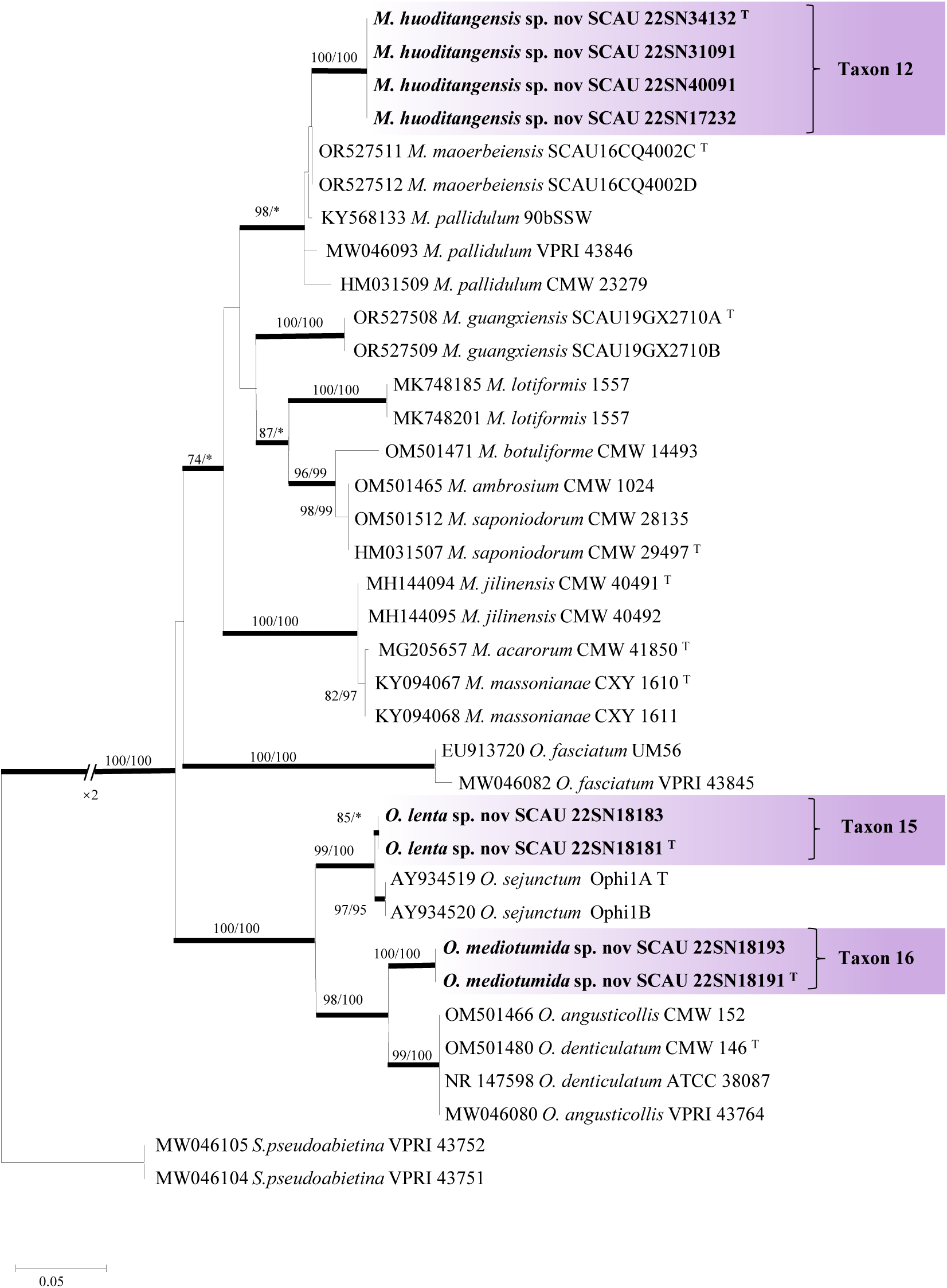
ML tree of *Masuyamyces* and“Group A”generated from the ITS sequence data. Sequences generated from this study are printed in bold. Bold branches indicate posterior probability values ≥0.95. Bootstrap values of ML/MP ≥ 70% are recorded at the nodes. T = ex-type isolates.

**Fig. 16.**
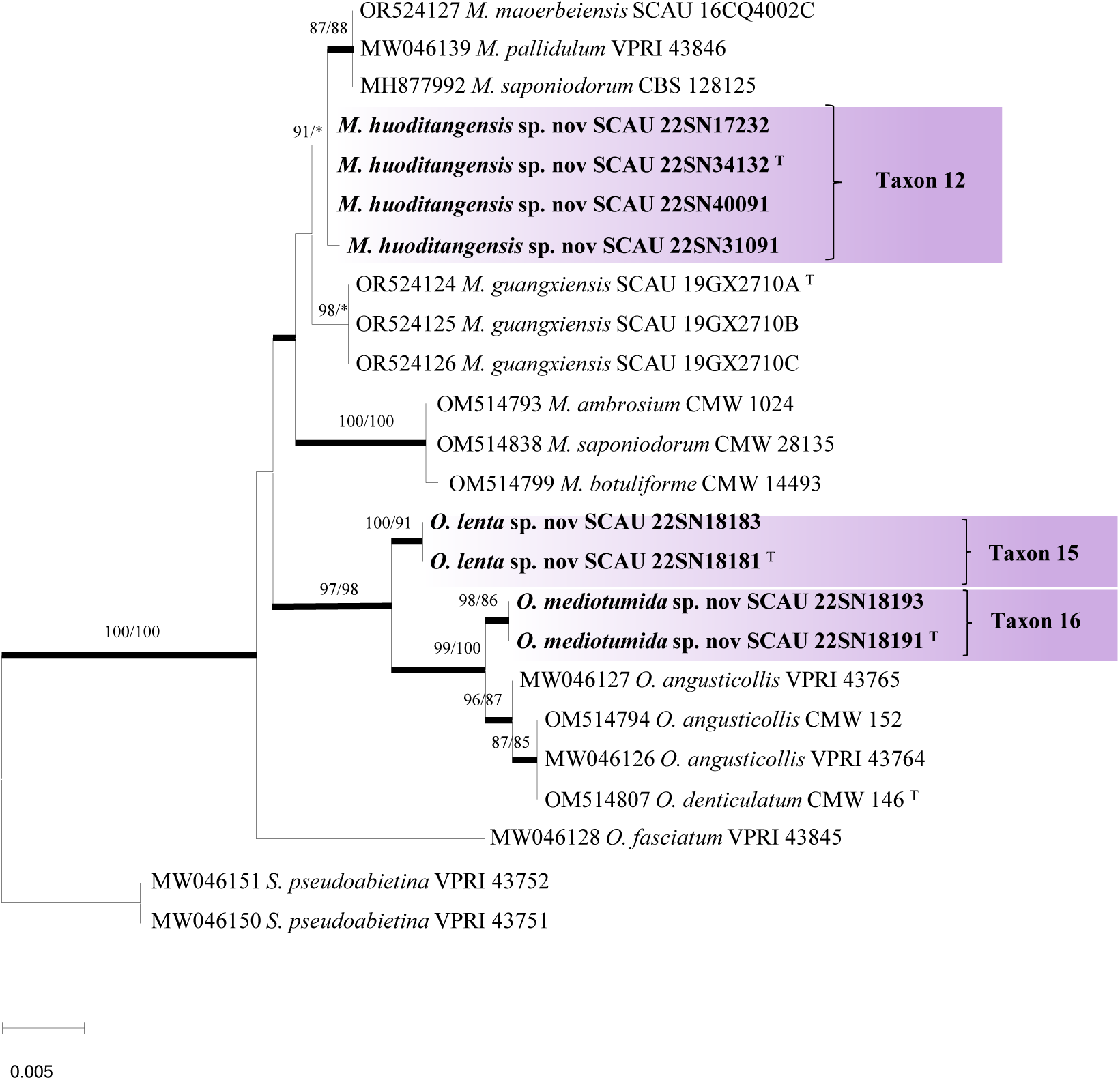
ML tree of *Masuyamyces* and“Group A”generated from the LSU sequence data. Sequences generated from this study are printed in bold. Bold branches indicate posterior probability values ≥0.95. Bootstrap values of ML/MP ≥ 70% are recorded at the nodes. T = ex-type isolates.

**Fig. 17.**
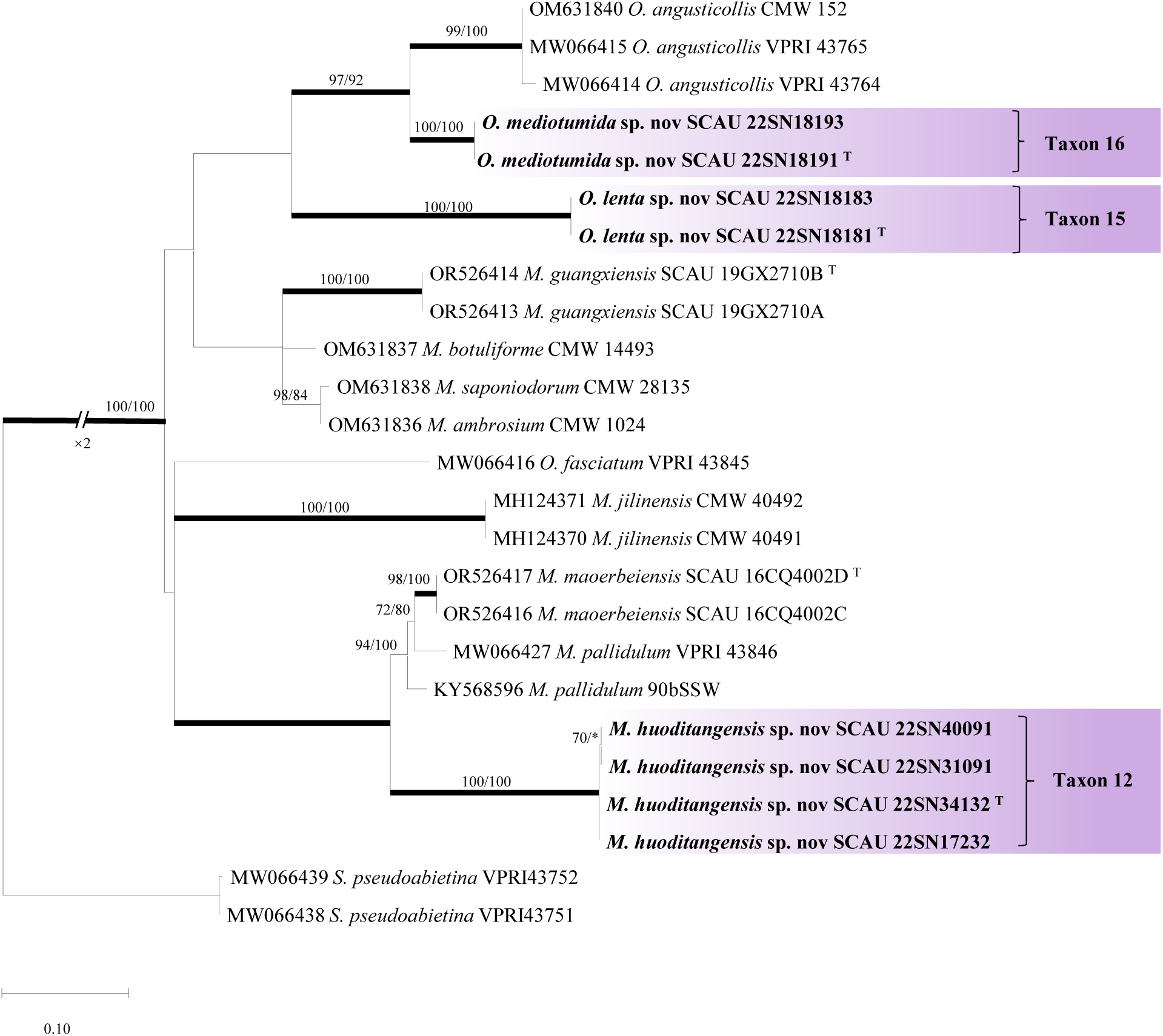
ML tree of *Masuyamyces* and“Group A”generated from the EF1-α sequence data. Sequences generated from this study are printed in bold. Bold branches indicate posterior probability values ≥0.95. Bootstrap values of ML/MP ≥ 70% are recorded at the nodes. T = ex-type isolates.

**Fig. 18.**
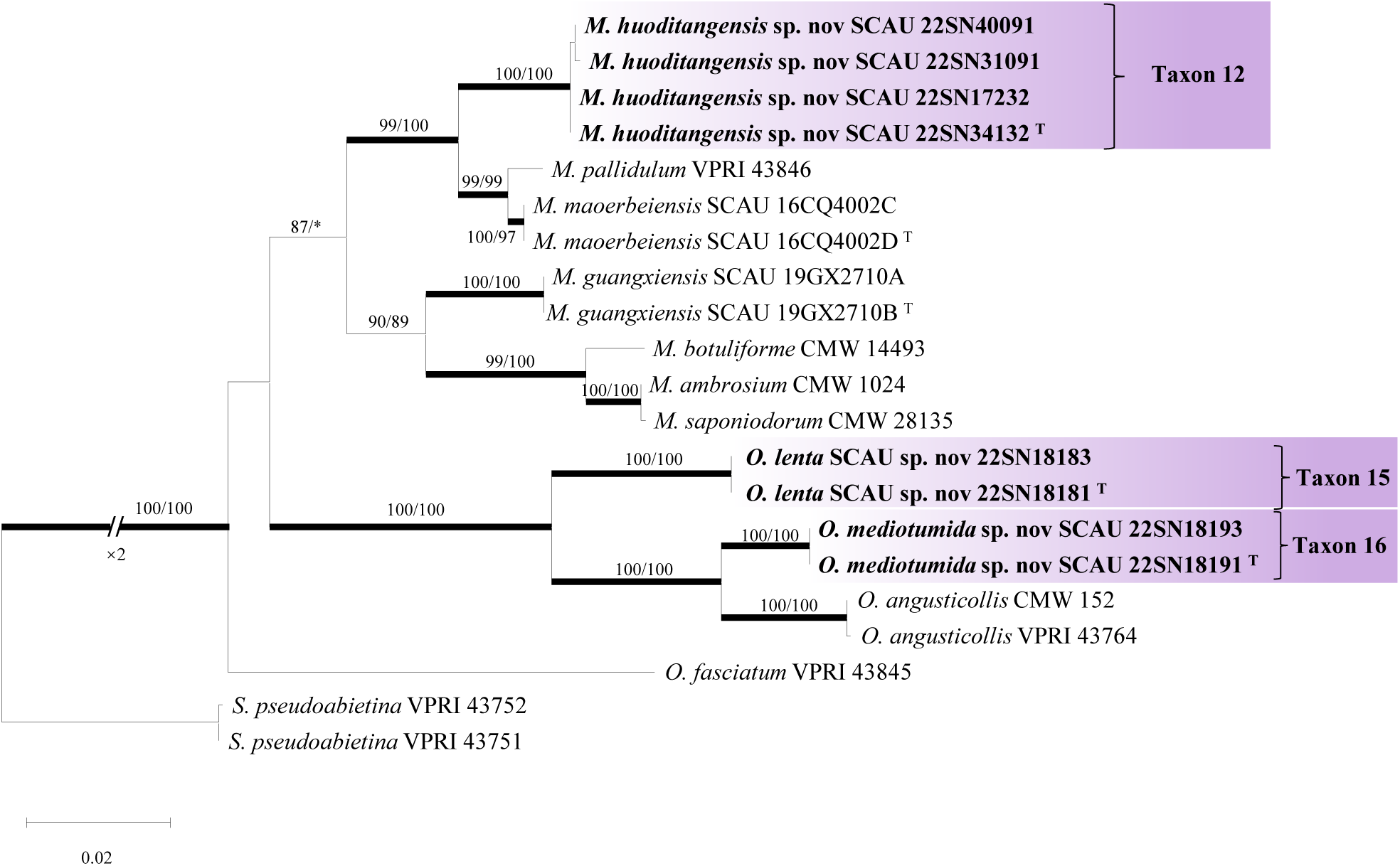
ML tree of *Masuyamyces* and“Group A”generated from the combined (ITS +ITS+ EF1-α) sequence data. Sequences generated from this study are printed in bold. Bold branches indicate posterior probability values ≥0.95. Bootstrap values of ML/MP ≥ 70% are recorded at the nodes. T = ex-type isolates.

The four representative strains of taxa 15 and taxa 16 belonged to “Group A”. According to the phylogenetic tree of βT, ITS, LSU, EF1-α and combined data set ( ITS + LSU + EF1-α ) ( Fig. 2, 15-18 ), taxa 15 and taxa 16 formed two branches with high node support values, which were closely related to *O. sejunctum* and *O. angusticollis*, respectively.

#### Ophiostoma clavatum complex

The ITS, LSU, EF1-α datasets and the j combined dataset ( βT + ITS + EF1-α ) of the *O. clavatum* complex are composed of 544, 779, 835 and 1725 characters, respectively, including gaps. According to the phylogenetic tree of βT, ITS, LSU, EF1-α and the combined dataset ( βT + ITS + EF1-α ) ( Fig. 2,19-22 ), the two representative strains of taxon 13 formed a branch with high node support, which was most closely related to *O. japonicum*. A representative strain of taxon 14 and a representative strain of taxon 17 were clustered on the same branch with *O. ips* and *O. minus* in the phylogenetic tree of BT ( Fig. 2 ) respectively, and had high node support. Therefore, taxon 14 and taxon 17 were determined to be previously discovered known species. Taxon 18 and 19 were clustered in the same branch with *O. shennongense* and *O. yaluense* in the phylogenetic trees of βT and ITS (Fig. 2, 19) with high node support values, respectively. Therefore, taxa 18 and 19 were identified as previously known species. Regrettably, both *O. shennongense* and *O. yaluense* are considered invalid names due to a lack of type specimens.

**Fig. 19.**
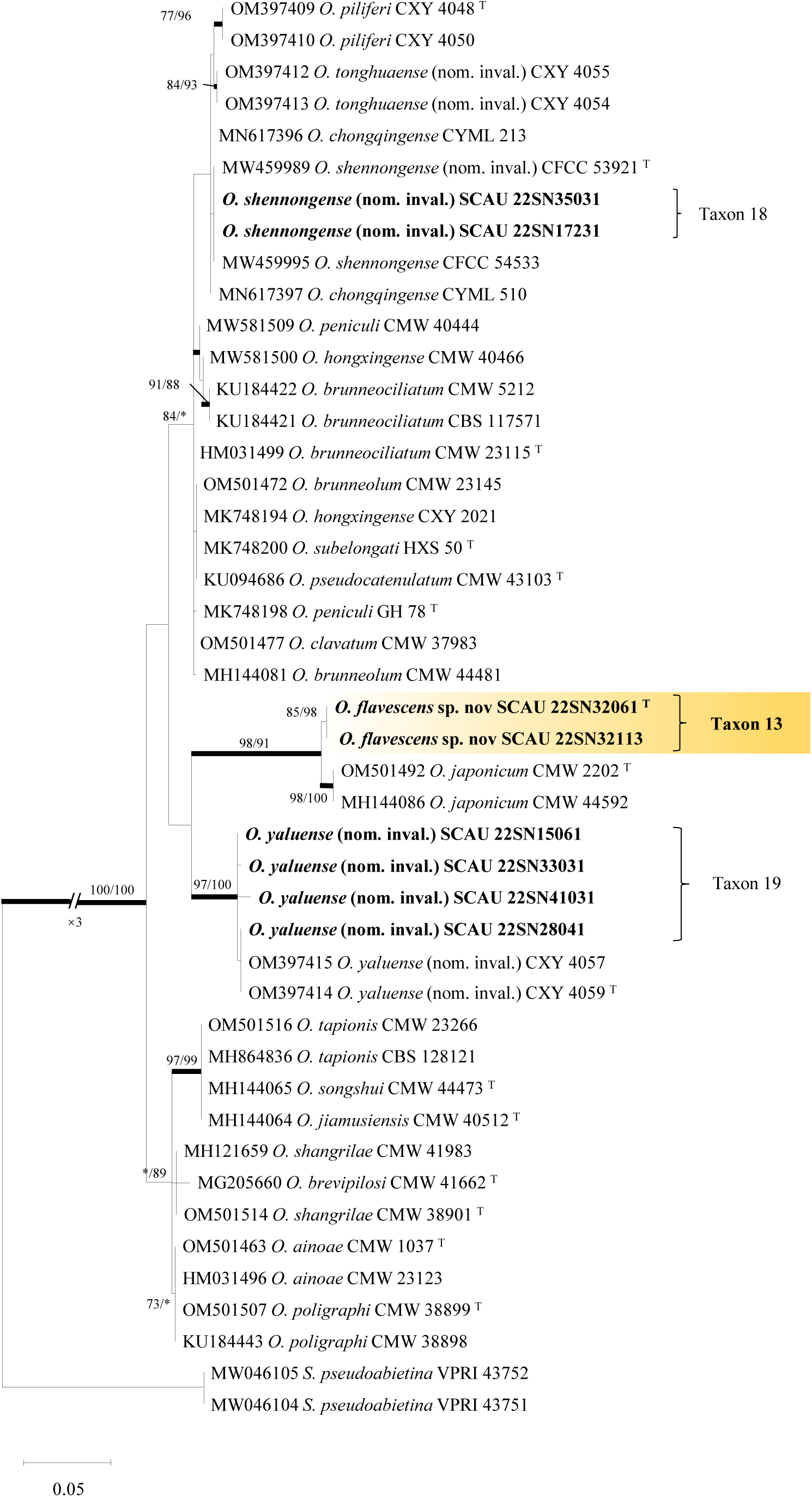
ML tree of *O. clavatum* complex generated from the ITS sequence data. Sequences generated from this study are printed in bold. Bold branches indicate posterior probability values ≥0.95. Bootstrap values of ML/MP ≥ 70% are recorded at the nodes. T = ex-type isolates.

**Fig. 20.**
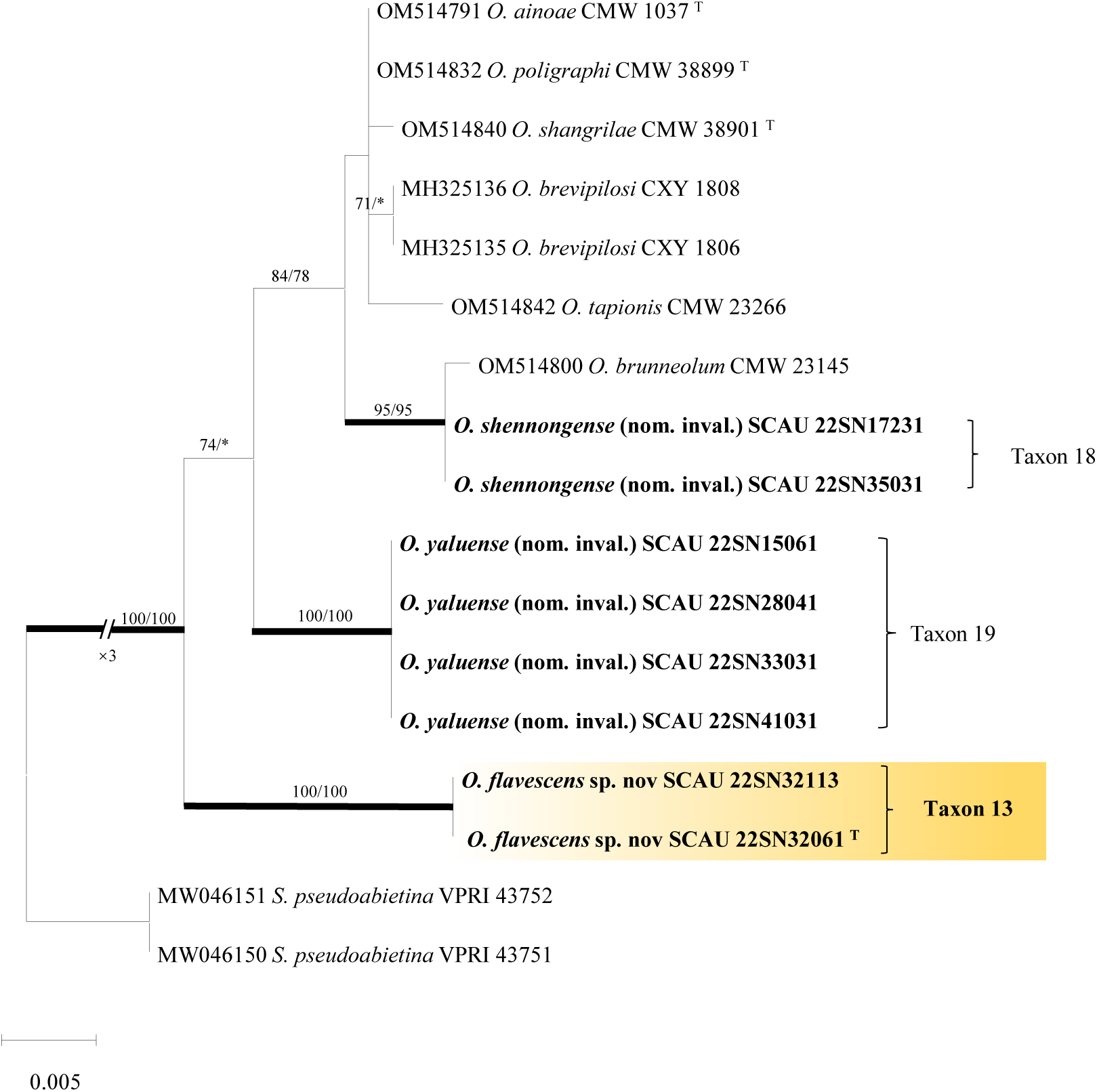
ML tree of *O. clavatum* complex generated from the LSU sequence data. Sequences generated from this study are printed in bold. Bold branches indicate posterior probability values ≥0.95. Bootstrap values of ML/MP ≥ 70% are recorded at the nodes. T = ex-type isolates.

**Fig. 21.**
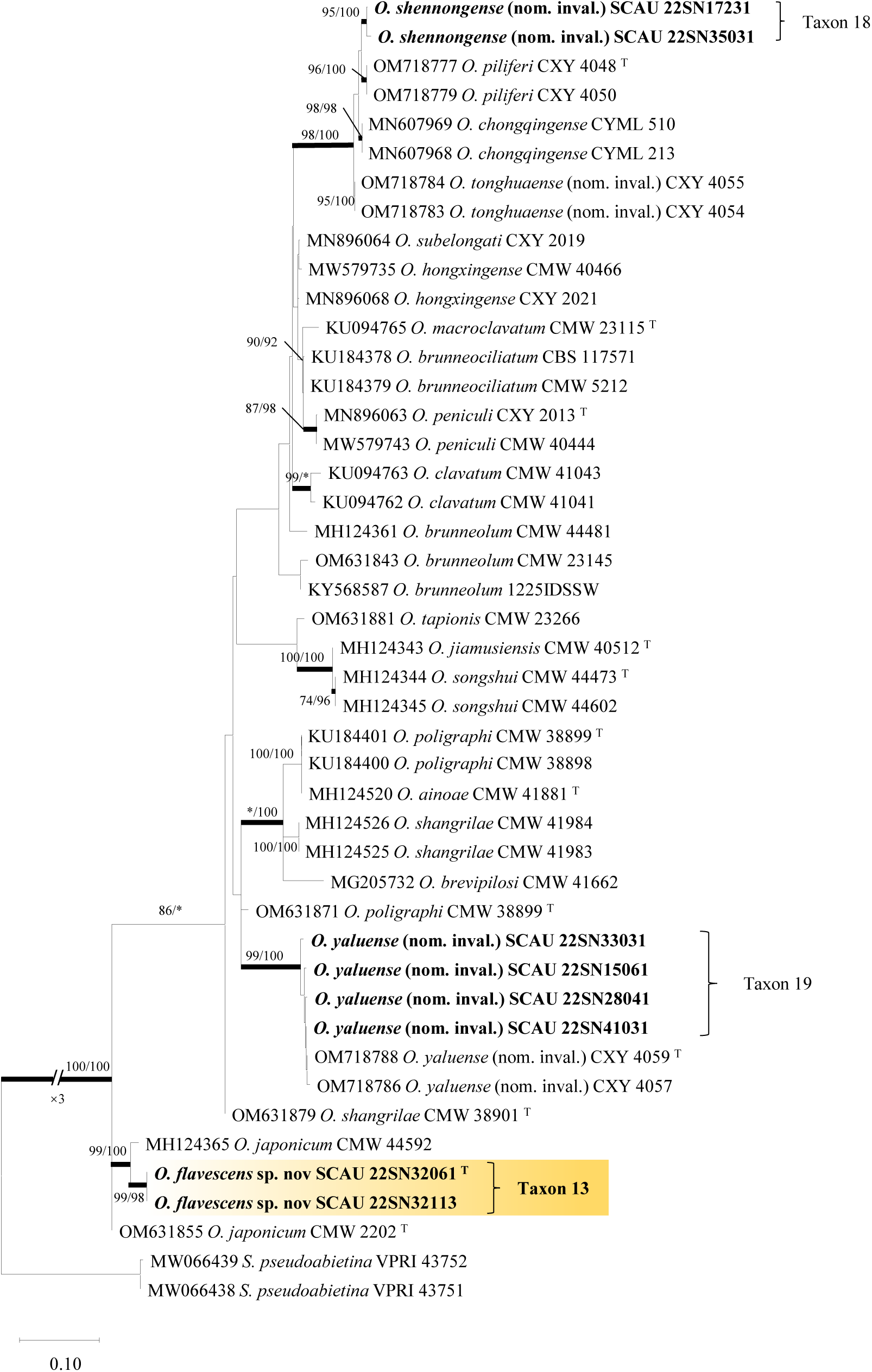
ML tree of *O. clavatum* complex generated EF1-α sequence data. Sequences generated from this study are printed in bold. Bold branches indicate posterior probability values ≥0.95. Bootstrap values of ML/MP ≥ 70% are recorded at the nodes. T = ex-type isolates.

**Fig. 22.**
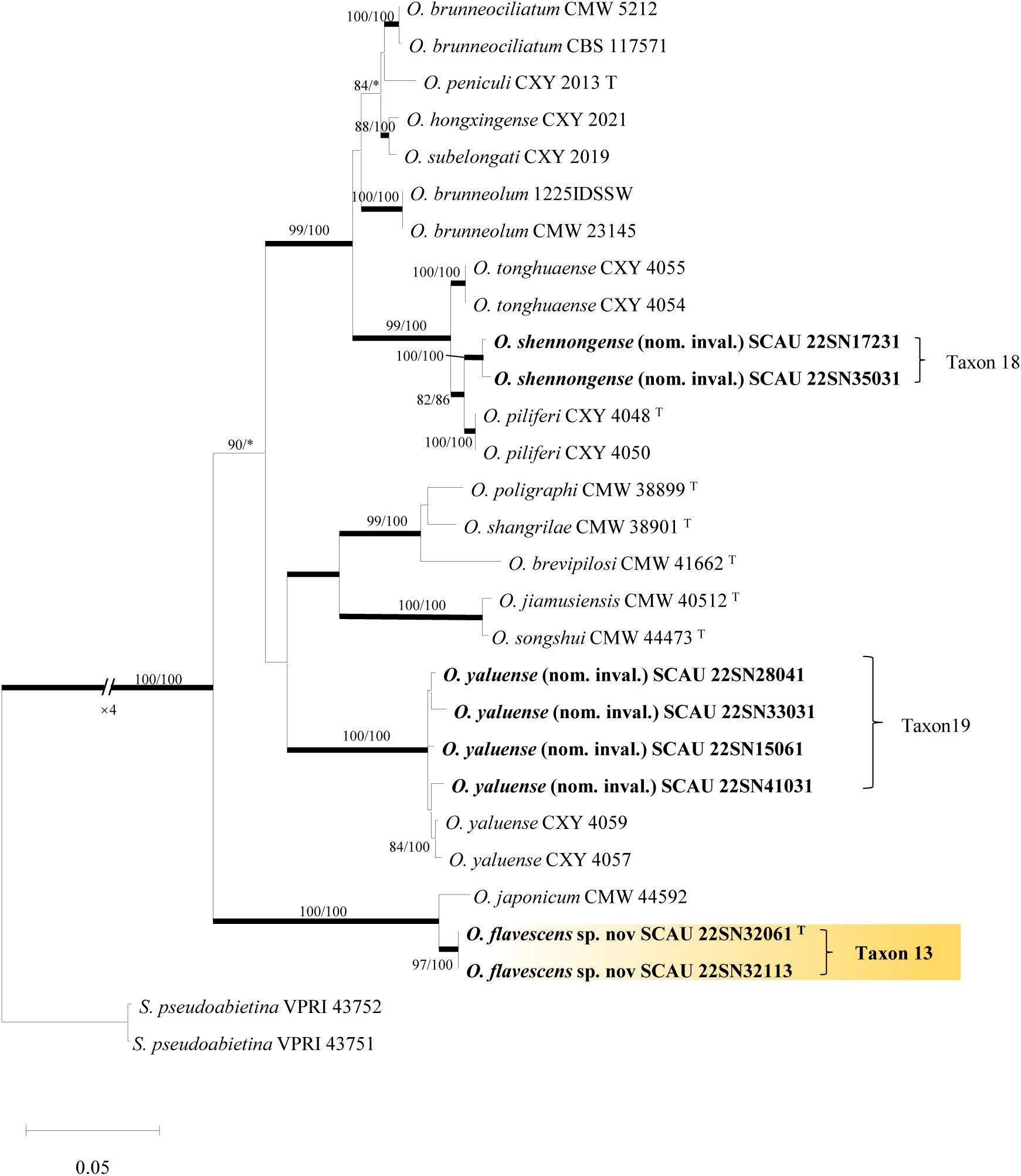
ML tree of *O. clavatum* complex generated from the combined (βT + ITS + EF1-α) sequence data. Sequences generated from this study are printed in bold. Bold branches indicate posterior probability values ≥0.95. Bootstrap values of ML/MP ≥ 70% are recorded at the nodes. T = ex-type isolates.

#### Sporothrix

Two representative strains of taxon 20 were clustered in the same branch with *S. pseudoabietina* in the phylogenetic tree of BT ( Fig. 2 ), so the taxon was defined as a previously discovered known species.

### TAXONOMY

Seven of the 20 taxa identified in this study were confirmed to represent different terminal clades and were interpreted as new species of Ophiostomatales.

*Ceratocystiopsis rugosa* M.J.Chen & M.L.Yin **sp. nov.**

MycoBank (Fig. 23)

**Fig. 23.**
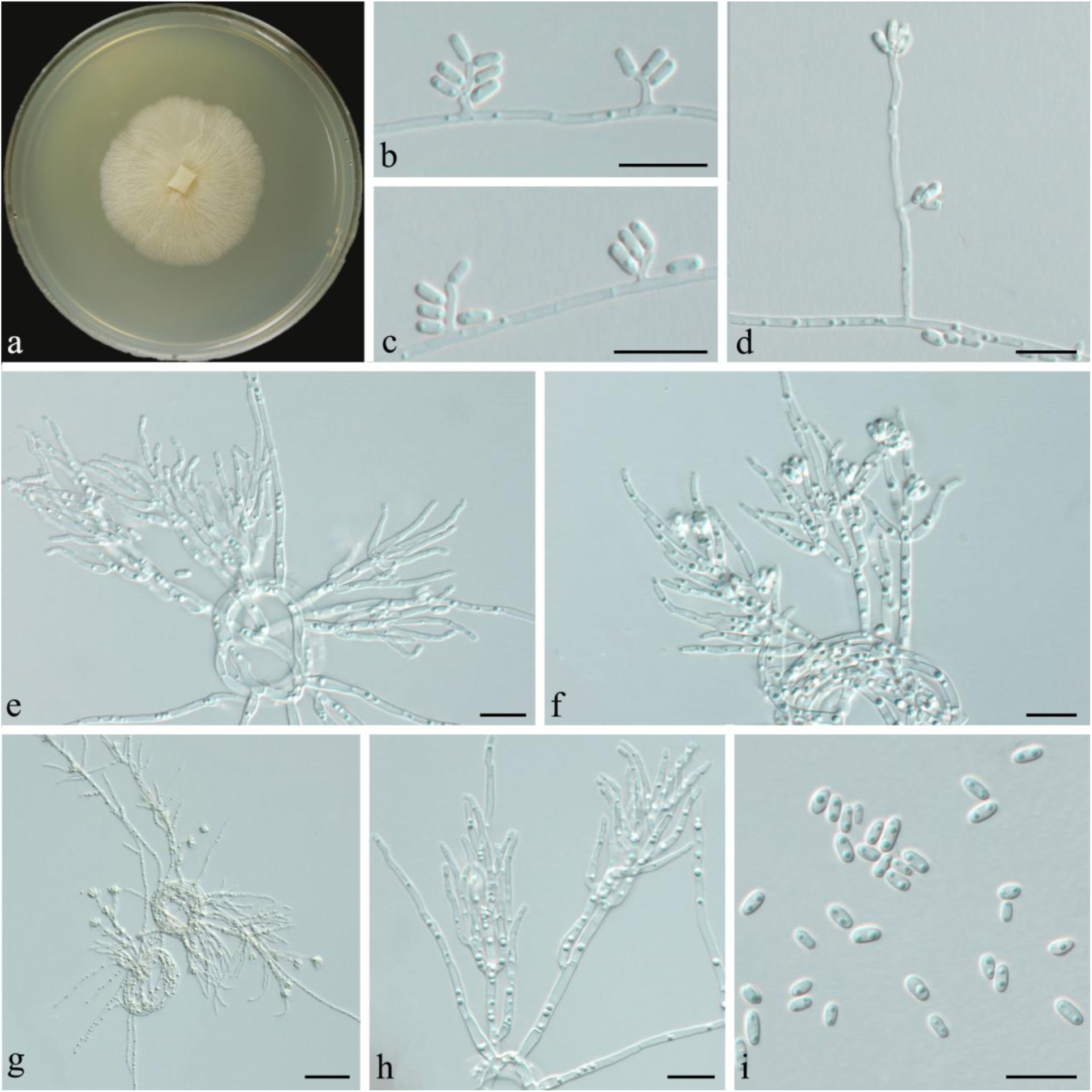
Morphological characteristics of ***Ceratocystiopsis rugosa*** (Taxon 2). a. Fourteen-d culture on 2%MEA. b-d. Simple and slightly branched micronematous conidiophores. e- h.branching macronematous conidiophores. i. Conidia of hyalorhinocladiella-like asexual morph. Scale bars: b-i=5μm.

*Etymology:* The epithet *rugosa* ( Latin ) refers to the wrinkled surface of the colony.

*Diagnosis: C. debeeria* is the closest phylogenetic relative of *C. rugosa.* The difference is that *C.rugosa* has small micronematous conidiophores and smaller conidia. In addition, the colony surface of *C.rugosa* was wrinkled, while that of *C.debeeria* was velvety.

*Type:* China, Shaanxi Province, Ankang City, 33 ° 43’N, 108 ° 45’E, collected from the gallery of *Polygraphus* sp. infesting *P. armandii*, Aug. 2022, Chen Minjie and Yin Mingliang (HAMS 352764-holotype; CGMCC 3.27261=SCAU 22SN25011 – ex-holotype culture).

*Description: Sexual morph* not observed.

*Asexual morph:* hyalorhinocladiella-like.

*Hyalorhinocladiella-like morph:* conidiophores micronematous or macronematous, abundantly formed. Micronematous conidiophores arising from vegetative hyphae are solitary, erect or slightly curved, transparent, usually producing 1-6 conidia, ( 1.5- ) 2.0-4.6 ( −6.4 ) × ( 0.4- ) 0.5-0.7 ( −0.8 ) μm. Macronematous conidiophores produced by coiled vegetative hyphae are erect, transparent, multibranched, ( 13.1- ) 18.2-27.6 ( − 33.6 ) × ( 0.9- ) −1.4 ( −1.6 ) μm; conidia transparent, smooth, oblong or elliptic, ( 1.6- ) 1.8-2.4 ( −2.7 ) × ( 0.9- ) −1.2 ( −1.3 ) μm.

*Culture characteristics:* Under the environmental conditions of 25 °C, *C. rugosa* grew on 2 % MEA medium for 14 d, and the radial diameter reached 41 mm. The growth rate was slow, the colony was yellow-white, the surface was rough and wrinkled, and the hyphae were attached to the surface of the medium without aerial hyphae. At 25 °C, the optimal growth rate was 2.9 ( ± 0.1 ) mm / d.

*Ecology:* Isolated from *Polygraphus* sp. infesting dying *P. armandii*.

Host trees: *P. armandii*.

*Distribution:* It is only known from Huoditang Forest Farm in Shaanxi Province. *Notes:* From a phylogenetic perspective, *C. rugosa* is closely related to *C. debeeria*. Although it is difficult to distinguish them based on LSU phylogenetic analysis, phylogenetic analysis of βT, ITS, and EF1-α shows that *C. rugosa* differs from *C. debeeria*. Moreover, the growth rate of the strain on MEA at 25 °C was 2.9 mm / d, which was faster than that of *C. debeeria* ( 0.82 mm / d ).

*Additional specimens examined:* China, Shaanxi Province, Ankang City, 33 ° 43’N, 108 ° 45’E, collected from the gallery of *Polygraphus* sp. infesting *P. armandii*, Aug. 2022, Chen Minjie and Yin Mingliang (SCAU 22SN25031= SCAU 22SN25111= SCAU 22SN25221= SCAU 22SN34141= SCAU 22SN41141).

***Graphilbum prolificum*** M.J.Chen & M.L.Yin **sp. Nov.**

MycoBank (Fig. 24)

**Fig. 24.**
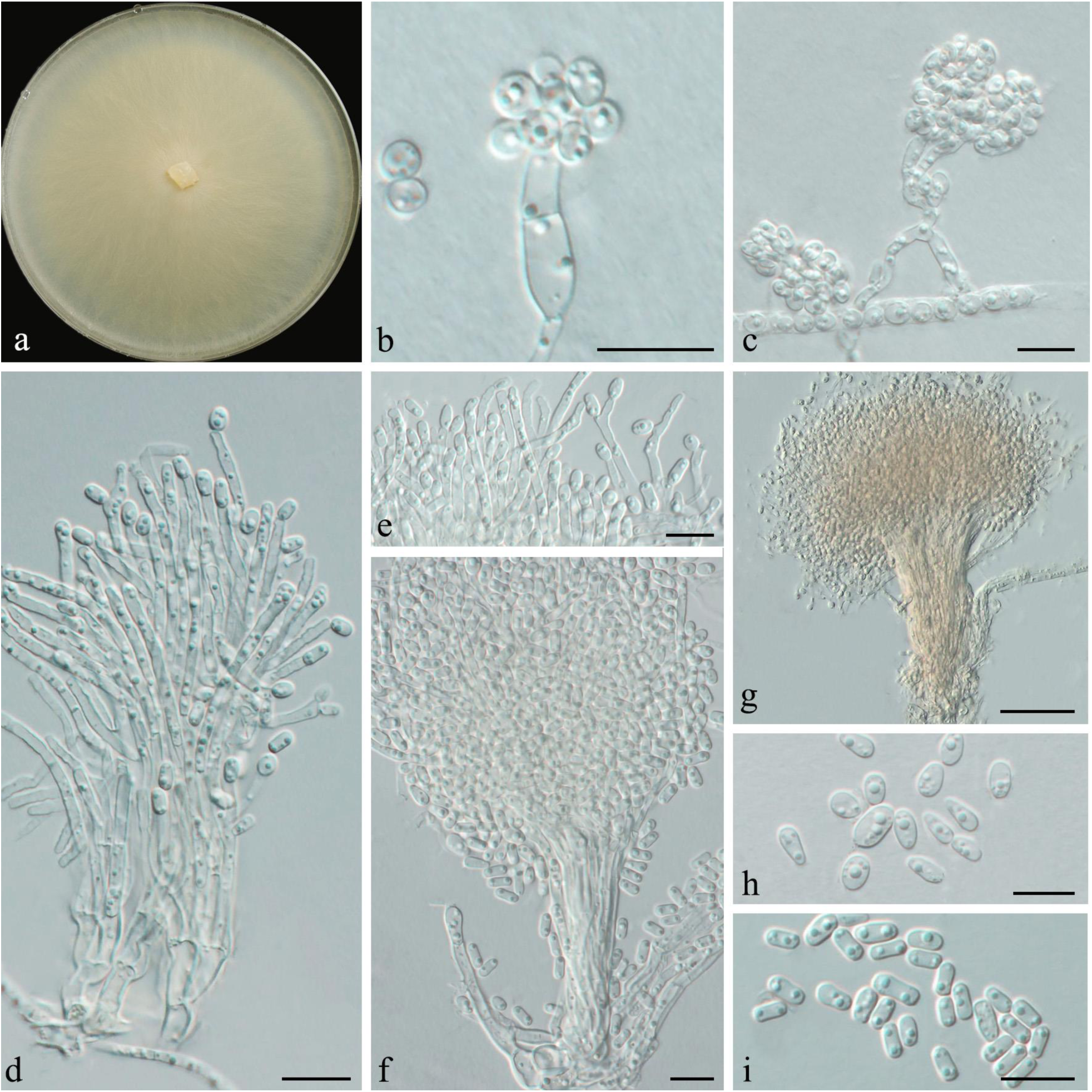
Morphological characteristics of ***Graphilbum prolificum*** (Taxon 6). a. Fourteen-d culture on 2%MEA. b-d. Hyalorhinocladiella-like asexual morph. e. Conidiogenous cells of pesotum-like macronematal asexual morph. f,g. Pesotum-like macronematal asexual morph. h. Conidia of hyalorhinocladiella-like asexual morph. i. Conidia of pesotum-like asexual morph. Scale bars: b,d-i=5μm,c=20μm.

*Etymology:* The epithet *prolificum* ( Latin ) refers to the prolific, this fungus thrives and has strong sporulation ability.

Diagnosis: The species was closely related to *G. crescericum* and *G. sexdentatum*, forming a lineage within *Graphilbum*. The phylogenetic analysis of ITS, βT, and EF1- α indicated that *G. prolificum* differed from *G. crescericum* and *G. sexdentatum*.

*Type*: China, Shaanxi Province, Ankang City, 33 ° 43’N, 108 ° 45’E, collected from the gallery of *Polygraphus* sp. infesting *P. armandii*, Aug. 2022, Chen Minjie and Yin Mingliang (HAMS 352765-holotype; CGMCC 3.27262=SCAU 22SN25051 – ex-holotype culture).

*Description*: *Sexual morph* not observed.

*Asexual morphs*: pesotum-like and hyalorhinocladiella-like.

*Pesotum-like morph:* macronematous, usually single, sometimes in groups, erect, translucent, rod-shaped, loosely arranged, growing on the surface of pine wood or medium, ( 64.2- ) 70.6-92.2 ( −94.6 ) μm long, including meristem, base width ( 4.7- ) 4.7-9.9 ( −12.0 ); conidiogenous cells were ( 9.2- ) 10.8-17.8 ( 21.8- ) × ( 0.9- ) 0.9-1.1 ( 1.2- ) μm, conidia are transparent, smooth, cylindrical to obovate, ( 2.0- ) 2.2-2.8 ( − 3.4 ) × ( 0.9- ) 0.9-1.3 ( −1.9 ) μm.

*Hyalorhinocladiella-like morph*: directly produced by mycelium submerged in the medium, Conidiophores often 2-3 branches, transparent, erect, ( 26.7- ) 28.8-49.2 ( − 47.5 ) × ( 1.5- ) 1.5-1.9 ( −2.0 ) μm ; conidia are transparent, smooth, aseptate, oval to cylindrical, occasionally concentrated at the top of the conidiophores, forming raspberry-like, ( 1.6- ) 1.8-2.2 ( −2.4 ) × ( 1.0- ) 1.1-1.3 ( −1.6 ) μm.

*Culture characteristics :* Under the condition of 25 °C, the radial diameter of *G. prolificum* reached 79 mm after 7 d of growth on 2 % MEA medium. The colony was white, with obvious texture, round shape and smooth edge. The mycelium was immersed in the medium with a small amount of aerial mycelium. The optimal growth was 11 ( ± 0.2 ) mm / d at 25 °C.

*Ecology:* Isolated from *Polygraphus* sp. infesting dying *P. armandii*.

Host trees: *P. armandii*.

*Distribution:* At present, it is only known from Huoditang Forest Farm in Shaanxi Province.

*Notes: G. prolificum* and *G. sexdentatum* have the closest genetic relationship in phylogenetic analysis. Both of them have pesotum-like and hyalorhinocladiella-like asexual structures. The difference is that *G. prolificum* produces smaller conidia than *G. sexdentatum.* Besides, at 25 °C, the growth rate of G. prolificum ( 11 mm / d ) on MEA was significantly faster than that of *G. sexdentatum* ( 5.1 mm / d ) ( P et al., 2014, Jankowiak et al., 2022 ).

*Additional specimens examined:* China, Shaanxi Province, Ankang City, 33 ° 43’N, 108 ° 45’E, collected from the gallery of *Polygraphus* sp. infesting *P. armandii*, Aug. 2022, Chen Minjie and Yin Mingliang ( SCAU 22SN24062=SCAU 22SN31031=SCAU 22SN33011=SCAU 22SN41171)。

***Graphilbum ramosus*** M.J.Chen & M.L.Yin **sp. Nov.**

MycoBank (Fig. 25)

**Fig. 25.**
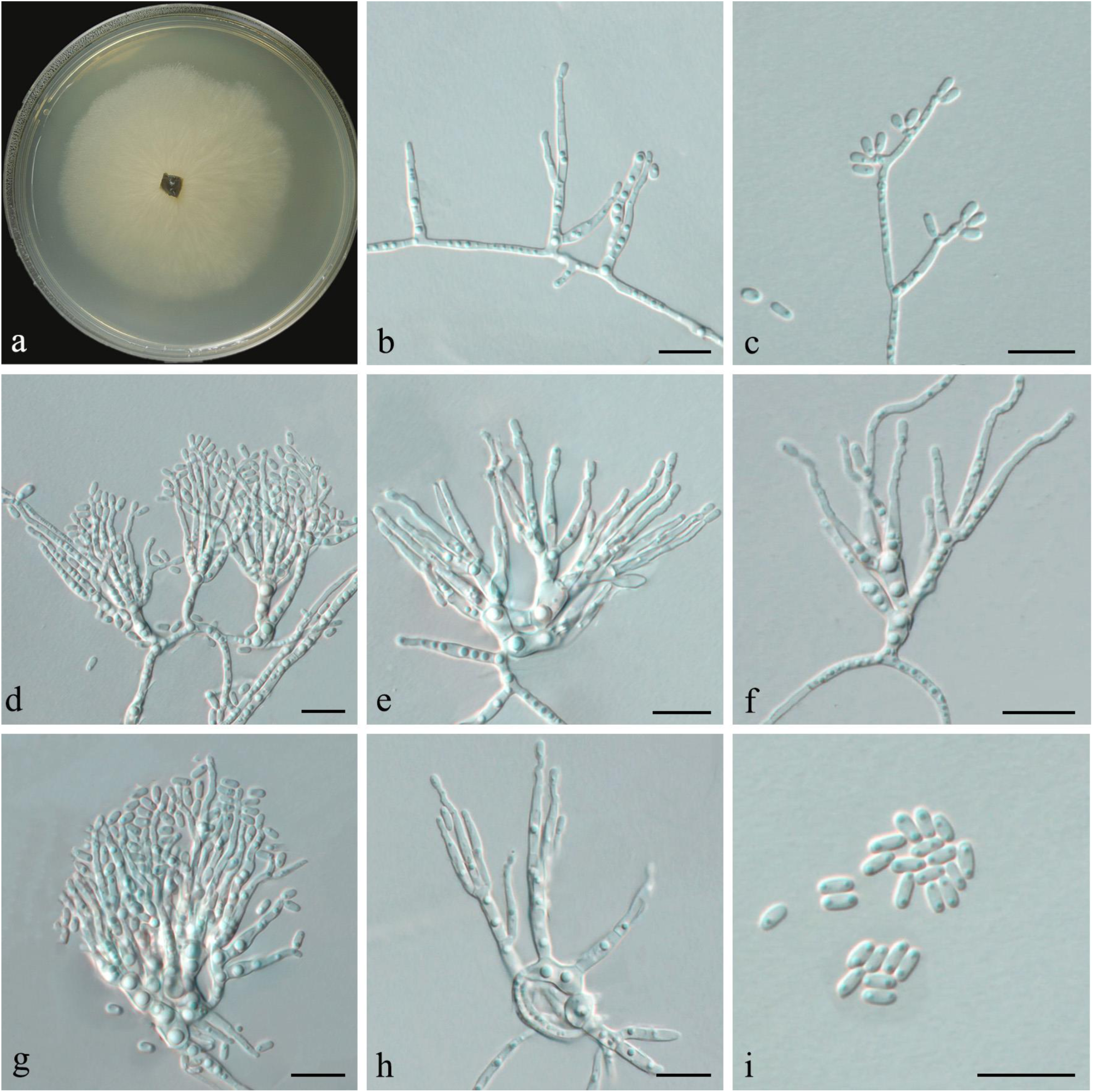
Morphological characteristics of ***Graphilbum ramosus*** (Taxon 7). **a.** Fourteen-d culture on 2%MEA. **b-h.** Hyalorhinocladiella-like asexual morph. **i.** Conidia of hyalorhinocladiella- like asexual morph. Scale bars: b-i=5μm.

*Etymology:* The epithet *ramosus* ( Latin ) refers to multi-branched or branched. The conidiophores of the fungus are often strongly branched to form a dendritic shape.

*Diagnosis:* Both *G. ramosus* and *G. gorcense* have hyalorhinocladiella-like conidiophores. The difference between them is that the conidiophores of *G. gorcense* are primarily solitary or occasionally branched, while the conidiophores of *G. ramosus* are usually accompanied by solid branches ( 5-8 branches ). Conidia of *G. license* ( 3- ) 3.5-4 ( −4.5 ) × ( 1- ) 1.5-1.5 ( −2 ) μm were more extensive than those of *G. ramosus* ( 1.6- ) 1.8-2.0 ( −2.1 ) × ( 0.8- ) 0.8- ( 1.1 ) μm. In addition, the growth rate of *G. gorcense* ( 5.9 mm / d ) on MEA at 25 °C was faster than that of *G. ramosus* ( 4.4 mm / d ).

*Type*: China, Shaanxi Province, Ankang City, 33 ° 43’N, 108 ° 45’E, collected from the gallery of *Dendroctonus armandi* infesting *P. armandii*, Aug. 2022, Chen Minjie and Yin Mingliang (HAMS 352766-holotype; CGMCC 3.27263=SCAU 22SN42141 – ex-holotype culture).

*Description*: *Sexual morph* not observed.

*Asexual morphs*: hyalorhinocladiella-like.

*Hyalorhinocladiella-like morph:* directly produced by mycelium submerged in agar, the conidiophores are transparent, the first layer is often accompanied by solid branches, forming a dendritic shape, ( 13.6- ) 15.9-22.3 ( −25.8 ) μm long, ( 1.0- ) 1.1-1.9 ( −2.3 ) μm wide at the base. Conidia are oblong, transparent, smooth, ( 1.6- ) 1.8-2.0 ( −2.1 ) × ( 0.8- ) 0.8- ( 1.1 ) μm.

*Culture characteristics:* Under the condition of 25 °C, the radial diameter of *G. ramosus* reached 61 mm after 14 d of growth on 2 % MEA medium. The colony was white and round, and the mycelium was immersed in the medium without aerial mycelium. After storage at four °C for some time, the color of the colony gradually changed from white to pure black. The optimal growth was 4.4 ( ± 0.1 ) mm / d at 25 °C.

*Ecology:* Isolated from *Dendroctonus armandi* infesting dying *P. armandii*.

Host trees: *P. armandii*.

*Distribution:* It is only known from Huoditang Forest Farm in Shaanxi Province. *Notes:* From the perspective of phylogeny, *G. ramosus* is closely related to *G. gorcense*, but the colony of *G. gorcense* will change from white to green-gray with age. No colony of *G. ramosus* was observed to change to green-gray, and an immature ascus was found in *G. license*, but no ascus was observed in *G. ramosus* ( Jankowiak et al., 2022 ).

*Additional specimens examined:* China, Shaanxi Province, Ankang City, 33 ° 43’N, 108 ° 45’E, collected from the gallery of *Dendroctonus armandi* infesting *P. armandii*, Aug. 2022, Chen Minjie and Yin Mingliang (SCAU 22SN42143).

***Masuyamyces huoditangensis*** M.J.Chen & M.L.Yin **sp. Nov.**

MycoBank (Fig. 26)

**Fig. 26.**
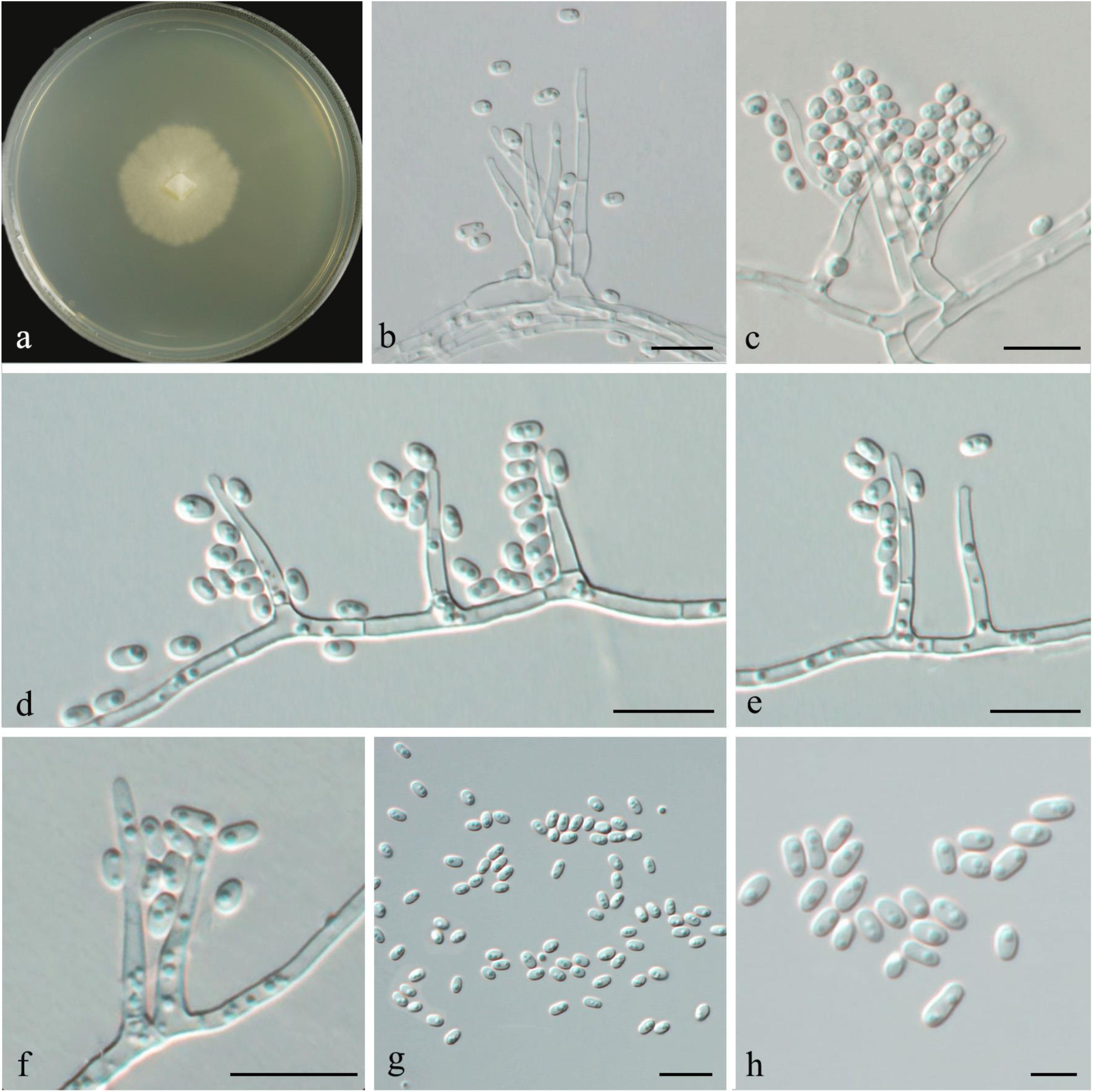
Morphological characteristics of ***Masuyamyces huoditangensis*** (Taxon 12). **a.** Fourteen-d culture on 2%MEA. **b-f.**Hyalorhinocladiella-like asexual morph. **g-h.** Conidia of hyalorhinocladiella-like asexual morph. Scale bars: b–h=5 μm.

*Etymology:* The epithet *huoditangensis* ( Latin ) refers to the forest farm of Huoditang, from which this fungus was collected.

Diagnosis: *M. huoditangensis* is closely related to *M. pallidulum*. The difference between them is that *M. huoditangensis* has smaller conidia.

*Type:* China, Shaanxi Province, Ankang City, 33 ° 43’N, 108 ° 45’E, collected from the gallery of *Dendroctonus armandi* infesting *P. armandii*, Aug. 2022, Chen Minjie and Yin Mingliang (HAMS 352767-holotype; CGMCC 3.27264=SCAU 22SN34132– ex- holotype culture).

*Description*: *Sexual morph* not observed.

*Asexual morphs*: hyalorhinocladiella-like.

*Hyalorhinocladiella-like morph:* conidiogenous cells arising directly from hyphae, transparent, septate, with a size of ( 5.1- ) 7.1-13.5 ( −17.3 ) × ( 0.9- ) −1.6 ( −2 ) μm. Conidia are transparent, smooth, ovoid to oblong, ( 1.3- ) 1.4-1.8 ( −2.1 ) × ( 0.9- ) 0.9-1.1 ( −1.2 ) μm in size.

*Culture characteristics:*Colonies on 2% MEA at 25 °C reaching 30 mm diam in14 d, the colony edge was smooth, the mycelium was attached to the medium, the growth was slow, and there was no aerial mycelium. The optimal growth was 2.1 ( ± 0.1 ) mm/ d at 25 °C.

*Ecology:* Isolated from *Dendroctonus armandi* infesting dying *P. armandii*.

Host trees: *P. armandii*.

*Distribution:* It is only known from Huoditang Forest Farm in Shaanxi Province. *Notes: M. huoditangensis* forms an independent branch in *Masuyamyces* and is closely related to *M. pallidulum*. Both species share a similar hyalorhinocladiella-like structure. The difference between the two is that the conidiophores and conidia produced by *M. huoditangensis* are smaller than those of *M. pallidulum*, and the growth rate of *M. huoditangensis* is slower than that of *M. pallidulum* at 25 °C. The colony growth rate of the former is 2 mm ( ± 0.1 ) mm / d, and the colony growth rate of the latter is 6 mm ( ± 0.1 ) mm / d.

*Additional specimens examined:* China, Shaanxi Province, Ankang City, 33 ° 43’N, 108 ° 45’E, collected from the gallery of *Dendroctonus armandi* infesting *P. armandii*, Aug. 2022, Chen Minjie and Yin Mingliang (SCAU 22SN17232=SCAU 22SN31091=SCAU 22SN40091).

***Ophiostoma flavescens*** M.J.Chen & M.L.Yin **sp. Nov.**

MycoBank (Fig. 27)

**Fig. 27.**
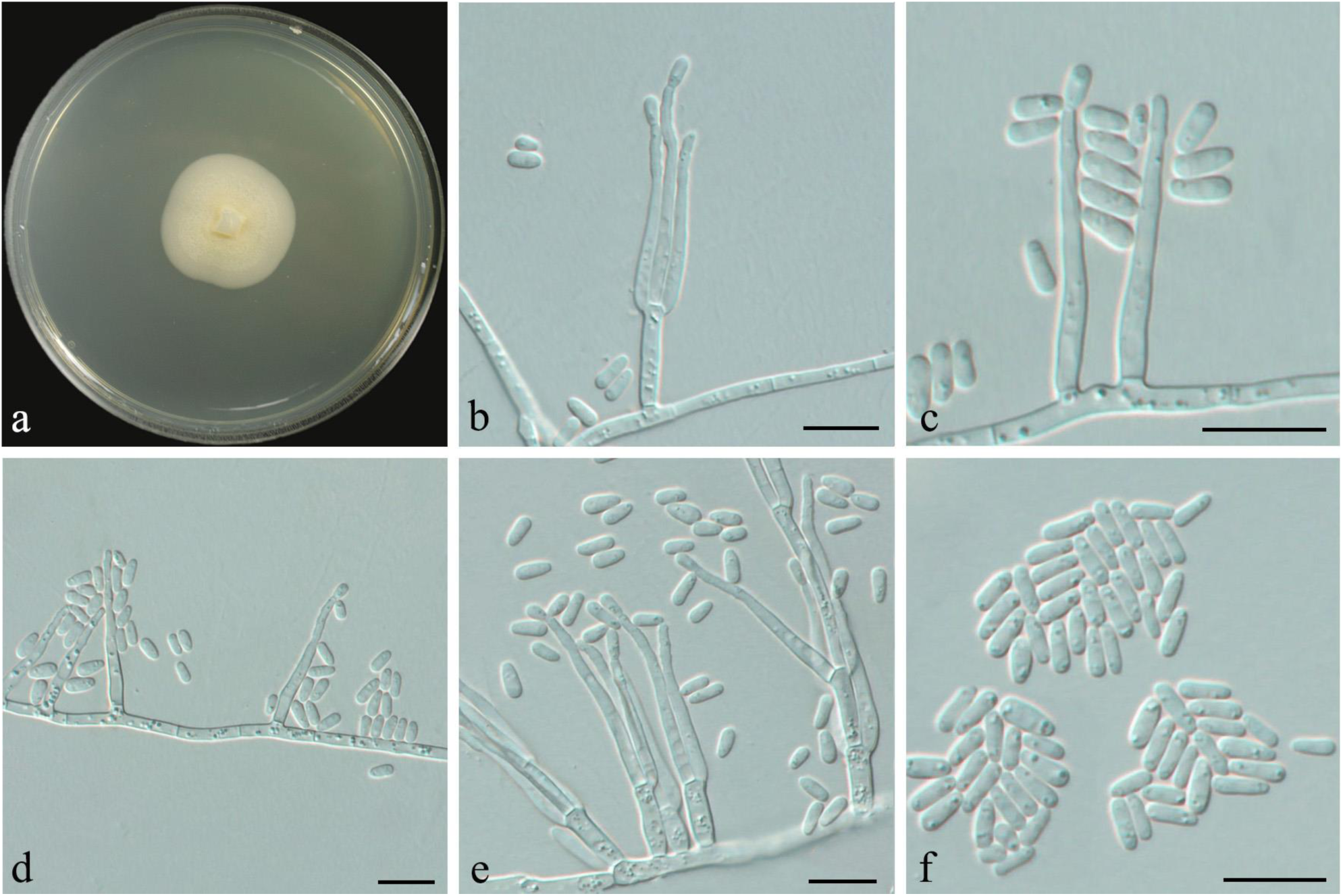
Morphological characteristics of ***Ophiostoma flavescens*** (Taxon 13). **a.** Fourteen-d culture on 2%MEA. **b-e.**Hyalorhinocladiella-like asexual morph. **f.** Conidia of hyalorhinocladiella-like asexual morph. Scale bars: b–f=5 μm.

*Etymology :* The epithet *flavescens* ( Latin ), which means that the color of the middle of the colony on MEA is light yellow.

*Diagnosis: O. flavescens* is closely related to *O. japonicum*. The difference between them is that *O. flavescens* does not have perithecium in vitro culture, and *O. primrose* has smaller conidia.

*Type*: China, Shaanxi Province, Ankang City, 33 ° 43’N, 108 ° 45’E, collected from the gallery of *Dendroctonus armandi* infesting *P. armandii*, Aug. 2022, Chen Minjie and Yin Mingliang (HAMS 352768-holotype; CGMCC 3.27265=SCAU 22SN32061– ex-holotype culture).

*Description*: *Sexual morph* not observed.

*Asexual morphs*: hyalorhinocladiella-like.

*Hyalorhinocladiella-like morph:* the conidiophores directly arise singly from the vegetative hyphae, or produce 1–3 branches, measuring ( 10.3- ) 14.0-25.0 ( −37.5 ) × ( 0.9- ) 1.1-1.5 ( −1.7 ) μm. Conidia transparent, smooth, cylindrical or clavate, ( 1.9- ) 2.4-3.0 ( −3.2 ) × ( 0.9- ) −1.2 ( −1.4 ) μm.

*Culture characteristics:*Colonies on 2% MEA at 25 °C reaching 29 mm diam. in 14 d, and the growth rate was slow. The inner ring of the colony was densely covered with light yellow villi, and the edge was smooth. The mycelium was immersed in the medium. The colony was light yellow, and the color gradually became lighter from the center to the periphery. With the prolongation of culture time, the local became black. The optimal growth rate was 2.0 ( ± 0.1 ) mm / d at 25 °C.

*Ecology:* Isolated from *Dendroctonus armandi* infesting dying *P. armandii*.

Host trees: *P. armandii*.

*Distribution:* At present, it is only known from Huoditang Forest Farm in Shaanxi Province.

*Notes: O. flavescens* and *O. japonicum* can be distinguished by phylogenetic analysis of the βT and EF1-α gene regions. In addition, the colony of *O. japonicum* primum albae, pars quidem coloniae viridifuscens, while the colony of *O. primrose* was light yellow at the beginning and turned black locally at the later stage ( YAMAOKA et al., 1997 ).

*Additional specimens examined:* China, Shaanxi Province, Ankang City, 33 ° 43’N, 108 ° 45’E, collected from the gallery of *Dendroctonus armandi* infesting *P. armandii*, Aug. 2022, Chen Minjie and Yin Mingliang (SCAU 22SN32113).

***Ophiostoma lenta*** M.J. Chen & M.L. Yin **sp. nov.** (Fig. 28)

**Fig. 28.**
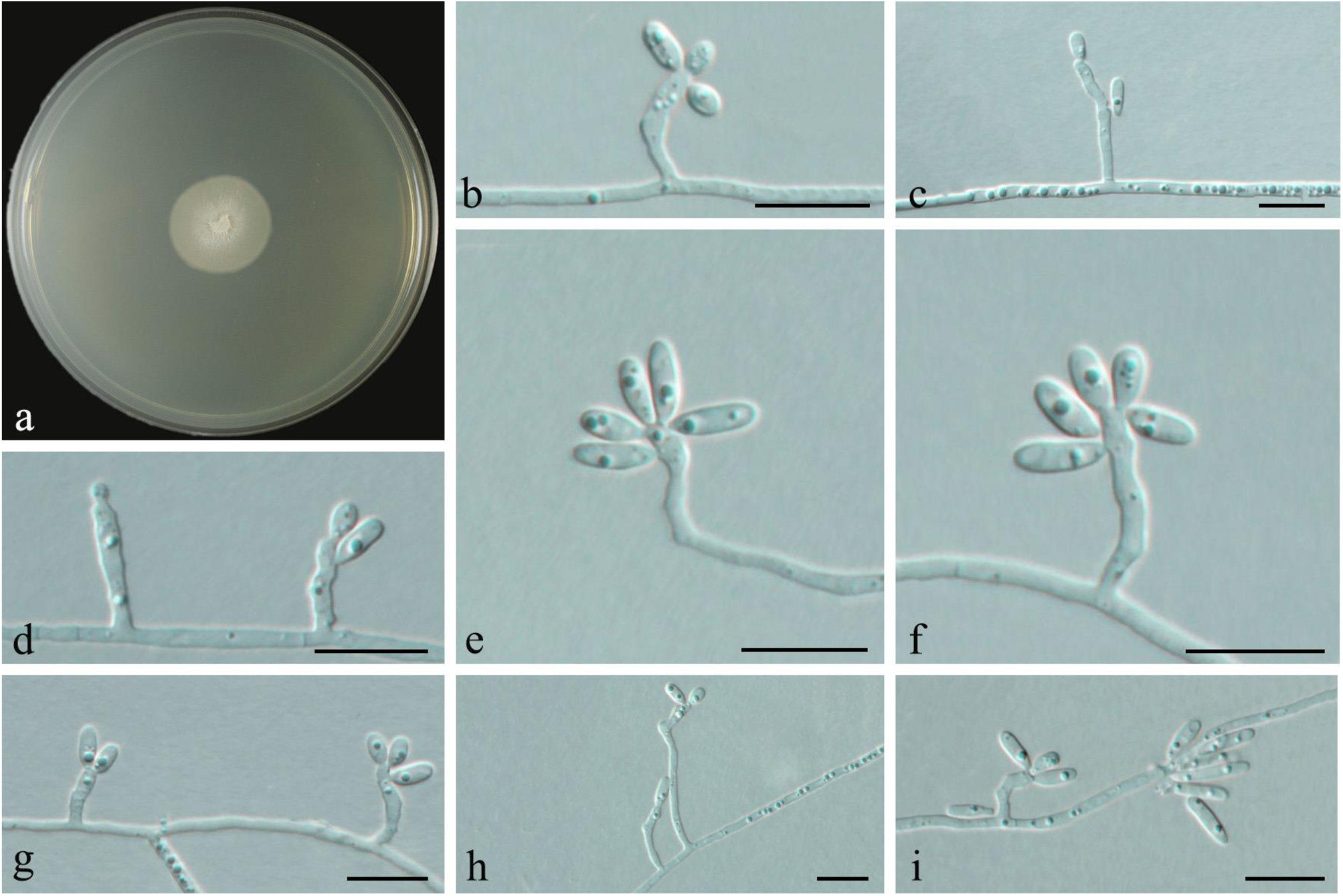
Morphological characteristics of ***Ophiostoma lenta*** (Taxon 15). **a.** Fourteen-d culture on 2%MEA. **b-i.** Hyalorhinocladiella-like asexual morph: conidiogenous cells and conidia. Scale bars: b–i=5 μm.

MycoBank XXXXXX

*Etymology:* The epithet *lenta*, refers to the slow growth rate of the strain.

*Type*: China, Shaanxi Province, Ankang City, collected from the gallery of *D. armandi* infesting *P. armandii*, Aug. 2022, Colls. M.J. Chen & M.L. Yin (**holotype** HAMS 352769; ex-holotype culture CGMCC 3.27266 = SCAU 22SN18181).

*Description*: Sexual state not observed. Asexual state hyalorhinocladiella-like. *Conidiophores* directly arising from the vegetative hyphae, conidiophores transparent, unbranched, straight or slightly curved, with rough edges, and the size is (2.5-) 3.8-9.2 (−14.3) × (0.6-) 0.7-0.9 (−1.1) μm. *Conidia* transparent, smooth, aseptate, clavate to ovate, (1.6-) 2.0 − 4 (−5.4) × (0.9-) −1.2 (−1.2) μm.

*Culture characteristics*: Colonies on 2 % MEA at 25 °C reached 18 mm diam. in 14 d. The colony was white, and the color gradually faded from the center to the periphery. The edge was smooth, with no aerial hyphae.

*Insect vector*: *Dryocoetes hectographus. Host trees*: *Pinus armandii*.

*Distribution*: Ankang, Shaanxi Province.

*Notes*: This fungus is closely related to *O. sejunctum*, but can be distinguished by the morphology of conidia. The conidia of *O. sejunctum* are obovate to ellipsoidal, whereas the conidia of *O. lenta* are clavate to ovate and are generally smaller in size. In addition,

*O. sejunctum* is characterized by the presence of allantoic ascospores and secondary and tertiary conidiophores, according to Villarreal et al. (2005). In contrast, *O. lenta* lacks these characteristics.

*Additional specimens examined:* China, Shaanxi Province, Ankang, from galleries of *Dendroctonus armandi* infesting *P. armandii*, Aug. 2022, Colls. M.J. Chen & M.L. Yin (culture SCAU 22SN18183).

***Ophiostoma mediotumida*** M.J.Chen & M.L.Yin **sp. Nov.**

MycoBank (Fig. 29)

**Fig. 29.**
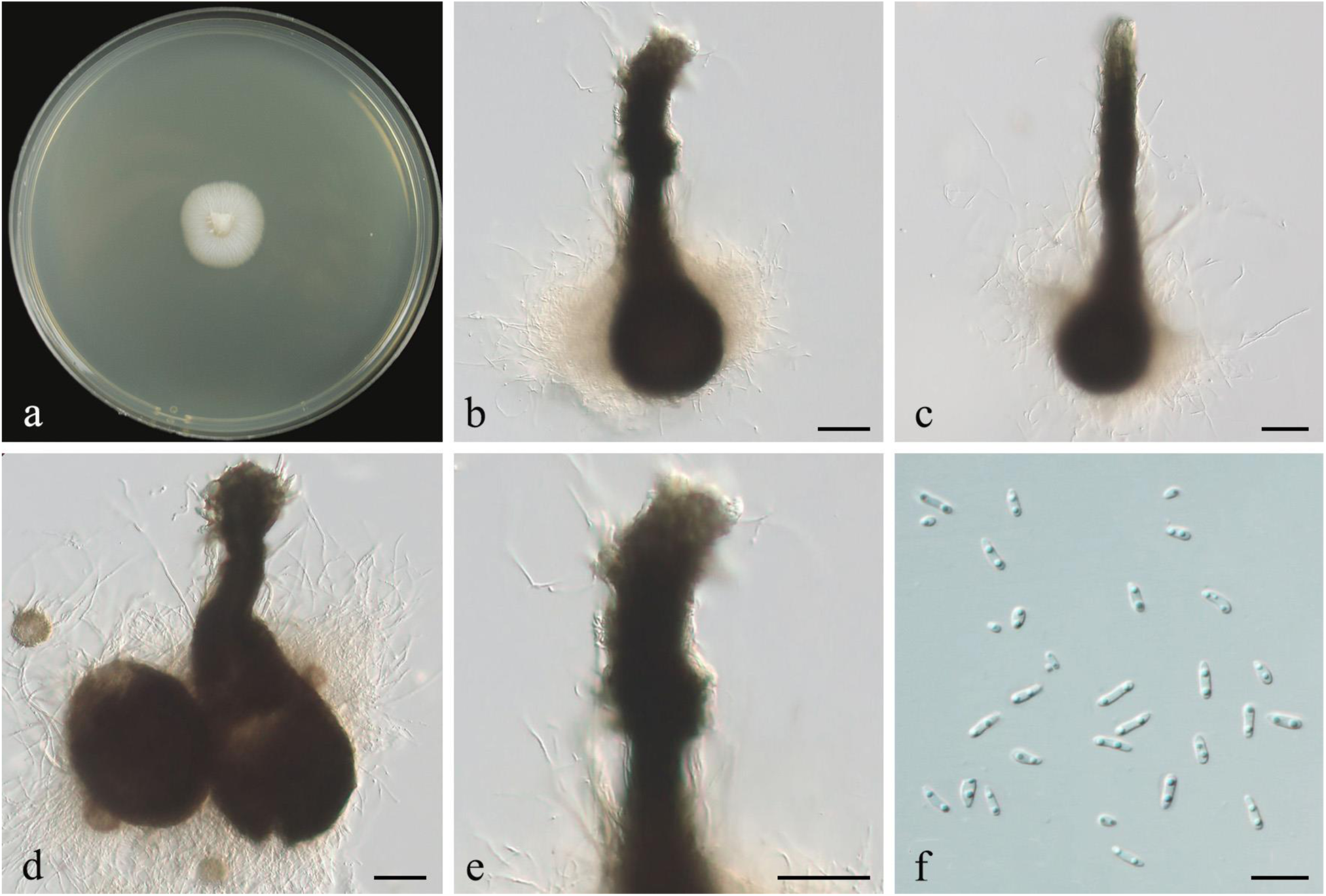
Morphological characteristics of ***Ophiostoma mediotumida*** (Taxon 16). **a.** Fourteen-d culture on 2%MEA. **b-d.** Perithecium. **e.** Top of neck. **f.** Ascospores. Scale bars: b-e=50 μm; f=5 μm.

*Etymology :* The epithet *mediotumida* ( Latin ), medio means middle ; tumida denotes a bulge, an enlarged part of the central part of the ascus.

*Diagnosis: O. mediotumida* is closely related to *O. angusticollis*. The difference is that *O. mediotumida* has a smaller ascus, the base of the ascus is pear-shaped or water-drop- shaped, the neck has a ring-shaped bulge, and the ascospores are long strips.

*Type*: China, Shaanxi Province, Ankang City, 33 ° 43’N, 108 ° 45’E, collected from the gallery of *Dendroctonus armandi* infesting *P. armandii*, Aug. 2022, Chen Minjie and Yin Mingliang (HAMS 352770-holotype; CGMCC 3.27267=SCAU 22SN18191– ex-holotype culture).

*Description*: *Sexual morph* perithecial. After 14 d, the perithecia appeared on the pine strip plate and grew on the surface of the agar or partially embedded in the agar. The base of the ascus shell was black, pear-shaped or water-drop-shaped, accompanied by transparent hyphae, with a diameter of ( 37.0- ) 40.7-49.5 ( −55.9 ) μm. Naturally extending out of the neck, straight or slightly curved, black, with a brush-like neck apex, accompanied by a ring-shaped bulge, full length ( 88.5- ) 113.6-156.0 ( −165.9 ) μm, no *ostiolar hyphae*. The basal width of the ascus neck is ( 13.8- ) 15.4-20.4 ( −23.7 ) μm, and the top width of the ascus neck is ( 10.0- ) 11.0-13.8 ( −14.7 ) μm. *Ascospores* hyaline, long striped, without sheath, aseptate, ( 1.7- ) 2.4-3.4 ( −4.0 ) × ( 0.6- ) 0.9-1.1 ( −1.2 ) μm.

*Asexual morphs* not observed.

*Culture characteristics:*Colonies on 2% MEA at 25 °C reaching 19 mm diam. in 14 d, milky white, the mycelium was dense, and the growth was adherent. The optimal growth rate was 1.3 ( ± 0.03 ) mm / d at 25 °C.

*Ecology:* Isolated from *Dryocoetes hectographus* infesting dying *P. armandii*.

Host trees: *P. armandii*.

*Distribution:* At present, it is only known from Huoditang Forest Farm in Shaanxi Province.

*Notes:* According to the phylogenetic tree, *O. mediotumida* belongs to the currently undefined “ GroupA ” and closely related to *O. angusticollis*. It can be well distinguished by phylogenetic analysis. Furthermore, compared to *O. angusticollis*, *O. mediotumida* does not have a transparent gelatinous cushion at the top of the ascospore neck, and ascospores are ejected from the top of the ascospore neck in a filamentous form ( Olchowecki et al. 1974 ).

*Additional specimens examined:* China, Shaanxi Province, Ankang City, 33 ° 43’N, 108 ° 45’E, collected from the gallery of *Dendroctonus armandi* infesting *P. armandii*, Aug. 2022, Chen Minjie and Yin Mingliang(SCAU 22SN18193).

## DISCUSSION

In this study, 225 strains of ophiostomatoid fungi were isolated from the gallery of *P. armandii* infected by bark beetles. Combined with morphological and multi-gene phylogenetic analysis ( βT, ITS, LSU, EF1-α ), 20 species of 6 genera were identified. They include seven previously undescribed species. We describe these new species as *C. rugosa*, *G. prolificum*, *G. ramosus*, *M. huoditangensis*, *O. flavescens*, *O. lenta* and *O. mediotumida*. In addition, four species with invalid nomenclature, *G. parakesiyea, L. qinlingense*, *O. shennongense* and *O. yaluense*, and ten known species were recorded. Among them, *C. armandii*, *C. weihaiensis*, *L. koraiensis* were first found in China, *G. fragrans*, *L. taigense*, *L. piceaperdum*, *O. ips*, *O. minus* and *S. pseudoabietina* were first discovered abroad. In our study, the dominant species were *O. shennongense*, followed by *O. ips*, *C. weihaiensis*, *G. parakesiyea*, and *S. pseudoabietina*, which accounted for 28.25 %, 11.66 %, 9.42 %, 9.42 %, and 9.42 % of the total number of ophiostomatoid fungi, respectively ( Table 2 ).

**Table 2.**
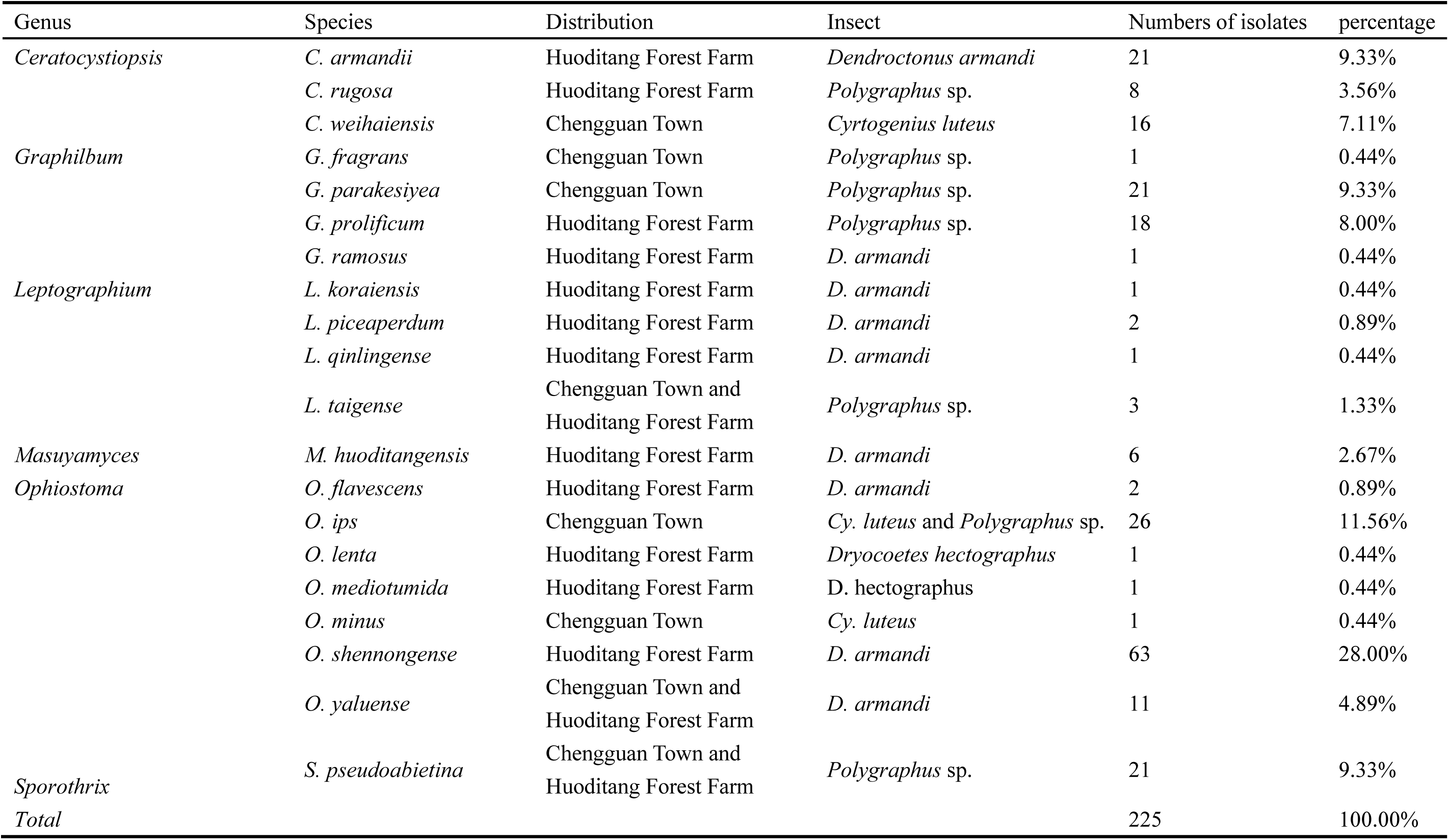
Strain numbers and percentage of various ophiostomatoid fungi isolated from Bark beetle galleries in western China.

In general, *O. shennongense* ( taxon 18 ) is the most common species in the ophiostomatoid fungi associated with bark beetles that infecting *P. armandii*. It is located in the *O. clavatum* complex ( Linnakoski et al. 2016 ).This species was previously discovered by Wang Huimin (Wang et al. 2022) and showed to be the dominant species in the ophiostomatoid fungi associated with *D. armandii*, but it was considered invalid due to the lack of model strains. This study confirmed the association between *O. shennongense* and *D. armandi* for the second time, and also indirectly indicated that there were a large number of *O. shennongense* in this area. The first ophiostomatoid fungi reported to be associated with *D. armandi* is *L. qinlingense*.In this study, only two strains were found. This species was first discovered by Tang et al. It has high pathogenicity and is a powerful assistant for *D. armandi* to invade *P. armandi* ( Pham et al. 2014 ). Tang et al. 2004 ; chen et al. 2004 ).

*Ceratocystiopsis* is widely distributed in Africa ( Nel et al. 2021 ), Asia ( Runlei et al. 2021 ) and Europe ( Jankowiak et al. 2021 ). At present, 34 species of the genus have been found, of which 4 species have been reported in China. In this study, a total of 45 strains of *Ceratocystiopsis* were isolated, among which *C. Weihaiensis* was first reported by Runlei et al., which was related to *Cryphalus piceae* ( Runlei et al. 2021 ). *C. rugosa* has the closest genetic relationship with *C. debeeria*, *C. debeeria* was first reported by Jankowiak et al., and it was proposed that *C. debeeria* is a specific exosymbiont of *P.chalcographii* ( Jankowiak et al. 2021 ). According to the size of the conidia and phylogenetic analysis, the two can be well distinguished.

According to phylogenetic analysis, *G. fragrans* ( taxon 4 ), *G. parakesiyea* ( taxon 5 ), *G. prolificum* ( taxon 6 ) and *G. ramosus* ( taxon 7 ) collected in this study belong to the genus *Graphilbum*, which is distributed in Africa ( Kamgan et al. 2008 ), Asia (Runlei et al. 2017; Min et al. 2019), Europe ( Pedro et al., 2014 ; jankowiak et al. 2020 ), South and North America ( Davidson, 1971; Geldenhuis et al. 2004; Reid et al. 2015 ), and has been extended to 28 formally described species. At present, no *Graphilbum* species have been reported as plant pathogens, and they are usually considered as humic organisms. They are mainly characterized by synnematous pesotum-like and mononematous hyalorhinocladiella-like asexual morphs ( Jankowiak et al. 2022 ). Taxon 6 is most closely related to the evolutionary branches composed of *G. sexdentatum*, *G. crescericum* and *G. furuicola*, and the morphological differences between them are very weak, which can be distinguished by the differences in molecular sequences. Taxon 7 was closely related to *G. gorcense*, which was found to be related to *Tetropium* sp. by Jankowiak et al. In *Graphilbum* genus, *G. gorcense* is the only species that does not form Synnemata and is the only species that produces gray-green colonies ( Jankowiak et al. 2020 ). Interestingly, no Synnemata structure was found in *G. ramosus* in this study. *G. Fragrans* was only one strain isolated in this study. It is very common in the genus *Graphilbum* and is thought to be composed of several mysterious species. For example, CBS 219.83 is considered to be the true source of *P. fragrans*, and then CBS 279.54 is considered to be a true version of the *P. fragrans* type (Harrington et al. 2001; Hafez et al. 2012; De Beer et al. 2013). *G. parakesiyea* was first discovered by Wang et al., which was proved to be associated with *D. armandi* ( Wang et al. 2022 ). In this study, 21 strains of *G. parakesiyea* were isolated from the beetle galleries of *P. armandii* infected by bark bettles, and the probability of separation was high, indicating that this species could be associated with different bark bettles.

*L. koraiensis* ( taxon 8 ) is located in the *L. galeiforme* complex ( Zhou et al. 2004 ), which is associated with the bark beetles of various attacking conifers ( kim et al. 2005; Linnakoski et al. 2012). *L. koraiensis* was first reported by Chang et al. ( Chang et al., 2019 ) to be related to the spruce bark beetle spruce bark beetle *Ips typographus*( Chang et al. 2019 ). In this study, only one *L. koraiensis* of this species was isolated, and it was related to *D. armandi*. *L. piceaperdum* ( taxon 9 ), first described by Jacobs et al. ( Jacobs et al. 2000 ), is located in the *L. piceiperdum* complex ( Linnakoski et al. 2012 ), in which all species have *leptographium*-like asexual morphology and cup-shaped ascospores ( Beer et al. 2013; Ando et al. 2015). In 2015, Ando et al. found that *L. piciceiperdum* was associated with a variety of bark beetles in Japan. Later, Chang et al. found that the species was associated with the *Ips typographus* in Jilin and Heilongjiang ( Ando et al. 2015; Chang et al. 2019 ). *L. taigense* ( taxon 11 ) was previously found to be associated with various conifer bark bettles in Russia ( Linnakoski et al. 2012 ). In 2017, Liu et al. discovered this species in Inner Mongolia, China, which was the first record in China ( Liu et al. 2017 ). Later, Runlei et al. were isolated from *Ips subelongatus* and *Ips typographus* in Inner Mongolia and Jilin, China ( Runlei et al. 2017 ; Chang et al. 2019 ). In Wang et al. study, it was proved that this species is weakly toxic to a variety of larch ( Wang et al. 2021 ), so we speculate that L.taigense is also weakly toxic to *P. armandii*, and can assist a variety of bark beetles in the area to attack *P. armandii*.

Taxon 12 is closely related to *M. pallidulum*, and they can be well distinguished by phylogenetic analysis. *M. pallidulum* was discovered by Linnakoski et al. ( Linnakoski et al. 2010 ) in pine and spruce trees in Finland, mainly related to hylastes brunneus erchison. Later, Jankowiak et al.showed that M.pallidulum could live as saprophytic bacteria in the roots of dead P.sylvestris, and had nothing to do with bark beetles that infect pine trees ( Jankowiak et al. 2012 ). Based on the genetic relationship between the two, we boldly speculate that taxon 12 has certain saprophytic ability.

*O. ips* ( taxa 14 ) belongs to the *O. ips* complex ( Beer et al. 2013 ). The species in this genus are characterized by cylindrical or allantoic ascospores with pillow-like sheaths and produce a series of asexual forms, which are described as Hyalorhinocladiella, *Leptographium* and *Pesotum*-like ( Zipfel et al. 2006 ; linnakoski et al. 2010 ), one of the most frequently isolated conchostomal fungi in China, is associated with various bark beetles that widely infect conifers ( Lu et al. 2009; Runlei et al. 2017 ; Wang et al. 2018Chang et al. 2019 ; Runlei et al. 2021 ), causing serious damage to conifer species in China. *O. flavescens* ( taxon 13 ) is closely related to *O. japonicum*. It was first discovered and reported to be related to *Ips typographus* in Japan ( Yamaoka et al. 1997 ).Later, Chang et al. also found and reported that *O. japonicum* was related to *Ips typographus* in Heilongjiang ( Chang et al. 2019 ). These two species can not only be distinguished by phylogenetic analysis, but also by morphological data. For example, *O. flavescens* does not have ascus in vitro culture and has smaller conidia.

The *O. lenta* ( taxon 15 ) and *O. mediootumida* ( taxon 16 ) in this study are currently in a smaller lineage and are not included in the currently recognized species complex commonly referred to as “Group A” ( Runlei et al. 2017; Wang et al. 2020 ; Trollip et al. 2021 ). Taxon 16 is closely related to *O. angusticollis*, which is related to *Dendroctonus adjunctus* in Spain ( Pérez Vera et al. 2011 ). Phylogenetic analysis of ITS showed that taxon 15 was closely related to *O. sejunctum*. It was known that *O. sejunctum* was from Spain and related to *Tomicus piniperda* on *P. pinaster* ( Villarreal et al. 2005 ). Since this species has only ITS sequence and lacks sequencing of other gene fragments, we distinguish it based on morphological differences. *O. sejunctum* is characterized by allantoic ascospores and the presence of secondary and tertiary conidiophores, but no sexual structure and secondary and tertiary conidiophores were found in taxon 15, only Hyalorhinocladiella-like conidiophores and a small number of long striped conidia were found.

In our study, we also recorded the known species *O. minus* ( taxon 17 ) in the *O. minus* complex ( Gorton et al. 2004 ), which is widely distributed and recorded in the Northern Hemisphere and China. It is one of the important fungi that cause wood blue stain (Gorton et al. 2000; Lu et al. 2009 ). Based on phylogenetic analysis and geographical differences, the *O. minus* lineage was divided into two exotic populations : the Eurasian branch and the North American branch. After that, Wang et al. divided *O. minus* into the third exotic population in the study of *Tomicus* species-related ophiostomatoid fungal community in southwestern China ( Wang et al. 2020 ; Gorton et al. 2004 ; Min et al. 2019 ). Different studies have shown that the Eurasian branch often combined with *Tomicus piniperda* or various bark beetles to kill endangered pine trees ( Gorton et al. 2000; Wang et al. 2020 ), while *O. minus* of the North American branch often interacts with *Dendroctonus frontalis* Zimmerman ( Barras et al. 1972 ).

In our study, one species belongs to the *S. gossypina* complex, *S. pseudoabietina* ( taxon 20 ), which is generally associated with various species of Cephalides and mites ( De Beer et al. 2016 ). This species was first discovered by Wang et al. in 2019 and isolated from the adults of *Tomicus yunnanensis* and *Tomicus minor*. Recently, the first record of *S. pseudoabietina* spread by *Xyleborus* was found in Australia. Studies have shown that this species has weak pathogenicity and does not show the potential to become the main pathogen of pine plants ( Mahony et al. 2023 ).

## CONCLUSIONS

In this study, we collected samples from Chengguan Town and Huoditang Forest Farm in Shaanxi Province. A total of 8 species of 5 genera were collected from Chengguan Town, and 15 species of 6 genera were collected from Huoditang Forest Farm. According to the phylogenetic data of the four DNA sequences, all the ophiostomatoid fungi obtained from Chengguan Town are known species, so we infer that the probability of finding new Ophiocranial fungi in this area in the subsequent investigation and separation is not high. In contrast, we found seven previously undiscovered species in the ophiostomatoid fungi collected from Huoditang Forest Farm, indicating that the ophiostomatoid fungi resources in this area have great potential for excavation.Although this study found many ophiostomatoid fungi in the galleries of bark beetles, the symbiotic mechanism between ophiostomatoid fungi and bark beetles and the pathogenicity of the new species still need to be further studied.

## Acknowledgments

We thank all the forest staff who assisted us during the sampling process in Guangdong, Fujian, Guangxi, Guizhou, Shandong, Liaoning and Shannxi Provinces over the years, and we especially appreciate Professor J. Wang for his unwavering support and care for our laboratory-related work.

## Declarations

### Availability of data and material

The manuscript mentioned all the availability of data. Ex-type cultures of new species were deposited in the Culture Collection of South China Agricultural University (SCAU) and the China General Microbiological Culture Collection Center (CGMCC). The type herbariums were preserved in the Fungarium (HAMS), Institute of Microbiology, Chinese Academy of Sciences. DNA sequence data are available in Genebank (https://www.ncbi.nlm.nih.gov/nucleotide/), and taxonomic novelties are available in Mycobank (https://www. mycobank.org).

### Competing Interests

The authors declare no conflicts of interest.

### Funding

This work was funded by the National Natural Science Foundation of China (32070012) and the Guangdong Basic and Applied Basic Research Foundation (2020A1515010486, 2022A1515010901).

### Authors’ contributions

Conceptualization, Mingliang Yin; Data curation, Minjie Chen; Funding acquisition, Mingliang Yin; Formal analysis, Minjie Chen; Funding acquisition, Mingliang Yin; Investigation, Kun Liu, Minjie Chen, Yutong Ran and Congwang Liu; Methodology, Mingliang Yin; Project administration, Mingliang Yin; Resources, Kun Liu, Minjie Chen, Yutong Ran and Congwang Liu; Software, Minjie Chen; Supervision, Mingliang Yin; Validation, Mingliang Yin; Visualization, Minjie Chen; Writing – original draft, Minjie Chen; Writing – review & editing, Tong Lin and Mingliang Yin.

## Notes

### Competing Interest Statement

The authors have declared no competing interest.

